# ATP2, the essential P4-ATPase of malaria parasites, catalyzes lipid-dependent ATP hydrolysis in complex with a Cdc50 β-subunit

**DOI:** 10.1101/2020.06.08.121152

**Authors:** Anaïs Lamy, Ewerton Macarini-Bruzaferro, Alex Perálvarez-Marín, Marc le Maire, José Luis Vázquez-Ibar

## Abstract

Efficient mechanisms of lipid transport are indispensable for the *Plasmodium* malaria parasite along the different stages of its intracellular life-cycle. Gene targeting approaches have recently revealed the irreplaceable role of the *Plasmodium-encoded* type 4 P-type ATPases (P4-ATPases or lipid flippases), ATP2, together with its potential involvement as antimalarial drug target. In eukaryotic membranes, P4-ATPases assure their asymmetric phospholipid distribution by translocating phospholipids from the outer to the inner leaflet. As ATP2 is a yet putative transporter, in this work we have used a recombinantly-produced *P. chabaudi* ATP2, PcATP2, to gain insights into the function and structural organization of this essential transporter. Our work demonstrates that PcATP2 heterodimerizes with two of the three *Plasmodium-encoded* Cdc50 proteins: PcCdc50B and PcCdc50A, indispensable partners for most P4-ATPases. Moreover, the purified PcATP2/PcCdc50B complex catalyses ATP hydrolysis in the presence of phospholipids containing either phosphatidylserine, phosphatidylethanolamine or phosphatidylcholine head groups, and that this activity is upregulated by phosphatidylinositol 4-phosphate. Overall, our work provides the first study of the function and quaternary organization of ATP2, a promising antimalarial drug target candidate.

## INTRODUCTION

Malaria is a global health problem affecting over 200 millions of people worldwide, and responsible in 2018 of nearly 405 000 deaths around the globe, mostly children (*World malaria report 2019*). Malaria is caused by parasites of the genus *Plasmodium*, a pathogen with a complex life-cycle between the transmission vector, the mosquito *Anopheles*, and a vertebrate host (Cowman et al. 2016). Five different *Plasmodium* species infect humans; *P. falciparum* being the most virulent one, accounting for ~90% of the mortality caused by this disease. Over the last 15 years, artemisinin-combined therapies (ACT) has permitted a substantial reduction of the mortality of this disease. However, *P. falciparum* strains resistant to current treatments, particularly to artemisinin, have already appeared in different regions of south Asia, representing a serious threat to eradicate malaria as no effective vaccine is yet available (Gosling and von Seidlein 2016). The fight against malaria thus requires a continuous supply of new drugs to supplement the existing therapeutic arsenal. In fact, the combination of drugs acting on different targets and/or stages is the most efficient strategy to combat malaria as it minimizes the development of new drug-resistance mechanisms (Burrows et al. 2017).

Recent efforts combining genetic disruption tools and phenotypic analysis in human and rodent malaria parasites have revealed that ~78% of *Plasmodium-encoded* membrane transport proteins (MTPs) are either essential for parasite’s survival or necessary for normal grow and development of the parasite (Kenthirapalan et al. 2016; Bushell et al. 2017; Zhang et al. 2018), thus confirming their therapeutic potential (Martin 2020). Among the annotated 144 MTPs, *P. falciparum* encodes 13 P-type ATPases (Martin 2020), most of them refractory to gene deletion (Weiner and Kooij 2016). P-type ATPases are a large family of primary transporters that utilize the energy from ATP hydrolysis to pump cations (subfamilies P1, P2 and P3), lipids (subfamily P4), or polyamine (P5 subfamily) (Palmgren and Nissen 2011; van Veen et al. 2020). In *Plasmodium*, P-type ATPases maintain ions homeostasis during its intracellular life-cycle (Kirk 2015), and some members are involved in drug research as the *P. falciparum* Na^+^/H^+^ P-type ATPase (PfATP4), the target of the antimalarial family of drugs spiroindolones (Rottmann et al. 2010; Rosling et al. 2018; Jiménez-Díaz et al. 2014). The *P. falciparum* homologue of the the sarco/endoplasmic reticulum Ca^2+^ ATPase (SERCA) pump (PfATP6) was initially proposed as an artemisinin target (Eckstein-Ludwig et al. 2003), although extensive work using recombinan PfATP6 did not support this (David-Bosne et al. 2016; Cardi et al. 2010). Recently, gene-targeting approaches have also uncovered the important role for parasite’s progression of the P4 subfamily, also known as lipid flippases (Bushell et al. 2017; Kenthirapalan et al. 2016; Zhang et al. 2018). In eukaryots, P4-ATPases maintains the asymmetric distribution of phospholipids in membranes by translocating phospholipids from the extracellular (or luminal) leaflet to the cytoplasmic one (Montigny et al. 2016; Andersen et al. 2016), key in important physiological functions as vesicle budding, coagulation or apoptosis (Xu et al. 2013; Paulusma and Elferink 2010). Moreover, new roles of these transporters have emerged in recent years as membrane lipids are also implicated in cell-signalling events (Sunshine and Iruela-Arispe 2017). In plants, the transport of lysophospholipids mediated by a P4-ATPase is involved in root development and size of stomatal apertures (Lisbeth R Poulsen et al. 2015). Also, the virulence of some pathogens and drug disponibility have been correlated with the expression and activity of P4-ATPases (Huang et al. 2016; Pérez-Victoria et al. 2006). The genome of *P. falciparum* contains four putative P4-ATPases: PfATP2, PfATP7, PfATP8 and PfATP11 (Martin 2020), well conserved in *Plasmodium* species with the exception of PfATP11 (Weiner and Kooij 2016). In addition, *P. falciparum* encodes two P4-ATPase-like proteins not present outside the Apicomplexa phylum: GCa and GCβ, consisting of a N-terminal P4-ATPase-like domain connected to a C-terminal guanylate cyclase domain. While there is some discrepancy with regard the vital role of ATP7 and ATP8 (Kenthirapalan et al. 2016; Bushell et al. 2017), all the studies agree that the activity of PfATP2 and its ortholog in the mouse malaria parasite *P. berguei* is completely irreplaceable during the blood stages. Moreover, in a recent study (Cowell et al. 2018) gene-duplication of the gene encoding PfATP2 was associated with drug-resistant phenotypes against two drugs belonging to the Malaria Box library of antimalarial compounds (Van Voorhis et al. 2016). This eventually situates PfATP2 as the target of these two antimalarial compounds or responsible of reducing the local concentration of these drugs at the target site.

P4-ATPases form heterodimeric complexes with members of the Cdc50/LEM3 protein family (Saito et al. 2004), and this association is essential for trafficking the P4-ATPase/Cdc50 complex, and for activity (Paulusma et al. 2008; Bryde et al. 2010). The recent Cryo-EM structures of two P4 ATPases, the *S. cerevisiae* Drs2p and the human ATP8A1, in complex with their respective Cdc50 subunits have revealed the structural basis of P4-ATPase/Cdc50 association (Timcenko et al. 2019; Hiraizumi et al. 2019). *P. falciparum* encodes three putative Cdc50 proteins (annotated as Cdc50A, Cdc50B and Cdc50C) also well conserved in *Plasmodium* species, and with eventual relevant roles for the parasite as judged by gene-disruption studies (Zhang et al. 2018). A recent study in *P. yoelii* has shown that Cdc50A interacts and stabilizes the GCβ/Cdc50A complex, mandatory for the gliding motility and midgut traversal of the ookinetes in the mosquito vector and, consequently, for parasite’s transmission (Gao et al. 2018). Also, its related Cdc50A homolog in *Toxoplasma gondii* is mandatory for bringing its GCβ partner to the plasma membrane prior cellular egression (Bisio et al. 2019). Despite the importance of P4-ATPase/Cdc50 association, to date, the identity of the Cdc50-interacting partners of ATP2 and the biological role of this association have not been studied, neither in the parasite nor in vitro using recombinant proteins.

ATP2 and the rest of *Plasmodium-encoded* P4-ATPases still remain as putative transporters waiting for functional annotation. Here, using recombinant *Plasmodium chabaudi* ATP2 (PcATP2), we have demonstrated that ATP2 associates with two *Plasmodium-encoded* Cdc50 proteins (PcCdc50B and PcCdc50A), and that the detergent-solubilized PcATP2/PcCdc50B complex catalyses ATP hydrolysis in the presence of phospholipid substrates. In addition, we have observed that phospholipid-activated ATP hydrolysis of PcATP2 is upregulated by PI4P. Overall, our work provides the first biochemical study of a *Plasmodium* P4-ATPase by assigning a functional annotation and by identifying two Cdc50-associated proteins.

## RESULTS

### 1. ATP2 and Cdc50 proteins are highly conserved in *Plasmodium* species and present in other apicomplexan parasites

The sequence alignments of ATP2 orthologs encoded by *P. falciparum*, PfATP2 (PF3D7_1219600), *P. vivax* (PVX_123625), and the malaria mouse models *P. berghei*, (PBANKA_1434800) and *P. chabaudi* (PCHAS_1436800) (Figure supplement 1) indicate a high degree of conservation (~60 % amino acid identity) of ATP2 among *Plasmodium*. In addition, P4-ATPases homologs to ATP2 are also present in the genome of other disease-causing intracellular parasites belonging, as *Plasmodium*, to the Apicomplexa phylum (Figure supplement 1) (Weiner and Kooij 2016), although it is still unknown if these transporters are also essential. These sequences contain the typical membrane topology of 10 predicted transmembrane domains (TMDs), the conserved P-type ATPase motif, DKTG, (intracellular loop between TMDs 4 and 5, where the initial aspartate is transiently autophosphorylated during the transport cycle), and the specific P4-ATPase motif DGET (intracellular loop between TMDs 2 and 3), implicated in protein dephosphorylation (Andersen et al. 2016). Interestingly, the glutamine in TMD1 and the two asparagine residues in TMDs 4 and 6, that coordinate the substrate’s phosphatidylserine (PS) moiety in ATP8A1 (Hiraizumi et al. 2019), are also conserved in these apicomplexan sequences (Figure supplement 1). Moreover, the well-conserved PICL (or PISL) motif present in TMD4 of P4-ATPases is also well conserved in the apicomplexan sequences (Figure supplement 1). In ATP8A1, the isoleucine and leucine residues of this motif also stablish hydrophobic contacts with the PS moiety of the substrate (Hiraizumi et al. 2019). This substrate-binding pocket present some similarities with the P2 subfamily of P-type ATPases like SERCA, where the isoleucine of this PICL motif is a glutamate that directly coordinates one of the two Ca^2+^ substrates (Olesen et al. 2007).

The genome of *P. falciparum* encodes three Cdc50 (or LEM) proteins: PfCdc50A (PlasmoDB ID: Pf3D7_0719500), PfCdc50B (PlasmoDB ID: Pf3D7_1133300) and PfCdc50C (PlasmoDB ID: Pf3D7_1029400). Like ATP2, each Cdc50 protein has close homologs in other *Plasmodium* species (50 to 60 % amino acid identity, see as example the alignments of Cdc50B orthologs in Figure supplement 2), as well as homologs in other apicomplexan parasites. All these sequences contain the two predicted TMDs connected by a large extracellular (or luminal) domain, as well as the four well-conserved cysteines, know to participate in two disulphide bridges (Timcenko et al. 2019; Hiraizumi et al. 2019). These disulphide bridges are important for the association of Cdc50 with the P4-ATPase and for the functional activity of the enzyme complex (Puts et al. 2012; Costa et al. 2016).

### 2. Screening co-expression and detergent solubilization of *Plasmodium* ATP2 and Cdc50 proteins

To unravel the functional and structural features of PfATP2, we examined its heterologous expression together with its putative Cdc50 proteins in the yeast *Saccharomyces cerevisiae* (Azouaoui et al. 2014; 2016). We first screened for expression of the ATP2 and Cdc50 sequences encoded by three different *Plasmodium* species: *P. falciparum, P. berghei* and *P. chabaudi* aiming at identifying an optimal ATP2 and Cdc50 candidate with suitable expression yield for biochemical studies. For western blot detection, we fused a biotin acceptor domain (BAD) at either the N-terminal (BAD-ATP2) or the C-terminal (ATP2-BAD) end of the ATP2 sequences, and a 10xHis-tag at the C-terminal end of each Cdc50 variant (Azouaoui et al. 2016). The *P. chabaudi* ortholog, PcATP2, was the only one that expressed in our system. ATP2 and Cdc50 sequences in *Plasmodium* species are highly conserved (~60 % amino acid identity), suggesting similar functional and structural features, thus making PcATP2 a fair paradigm of PfATP2.

Figure 1 shows the expression profiles of the two BAD-tagged versions of PcATP2, alone, or co-expressed with each of the three *P. chabaudi-encoded* Cdc50 proteins. PcATP2 displayed a slightly different electrophoretic mobility depending on the position of the BAD tag (Figure 1), previously observed in other P4-ATPase tagged as well with the BAD at both extremities (Azouaoui et al. 2014). Also, BAD-PcATP2 displayed two electrophoretic gel-bands, being the fastest one the most intense, indicating either a cleavage near the C-terminal end or different electrophoretic mobilities (Figure 1, lanes 2, 4, 6 and 8). Western-blot analysis also revealed that the three putative PcCdc50 proteins were also co-expressed with both BAD-PcATP2 and PcATP2-BAD (Figure 1, bottom panels). PcCdc50A displayed two electrophoretic mobilities, suggesting protein glycosylation events, typical of Cdc50 proteins (Figure 1, lanes 7 and 8). Overall, when either BAD-PcATP2 or PcATP2-BAD were co-expressed with each of the PcCdc50 proteins neither the electrophoretic pattern nor the intensity of the PcATP2 bands changed substantially. We discarded further studies with PcCdc50C because of its very low expression yield.

**Figure 1.**
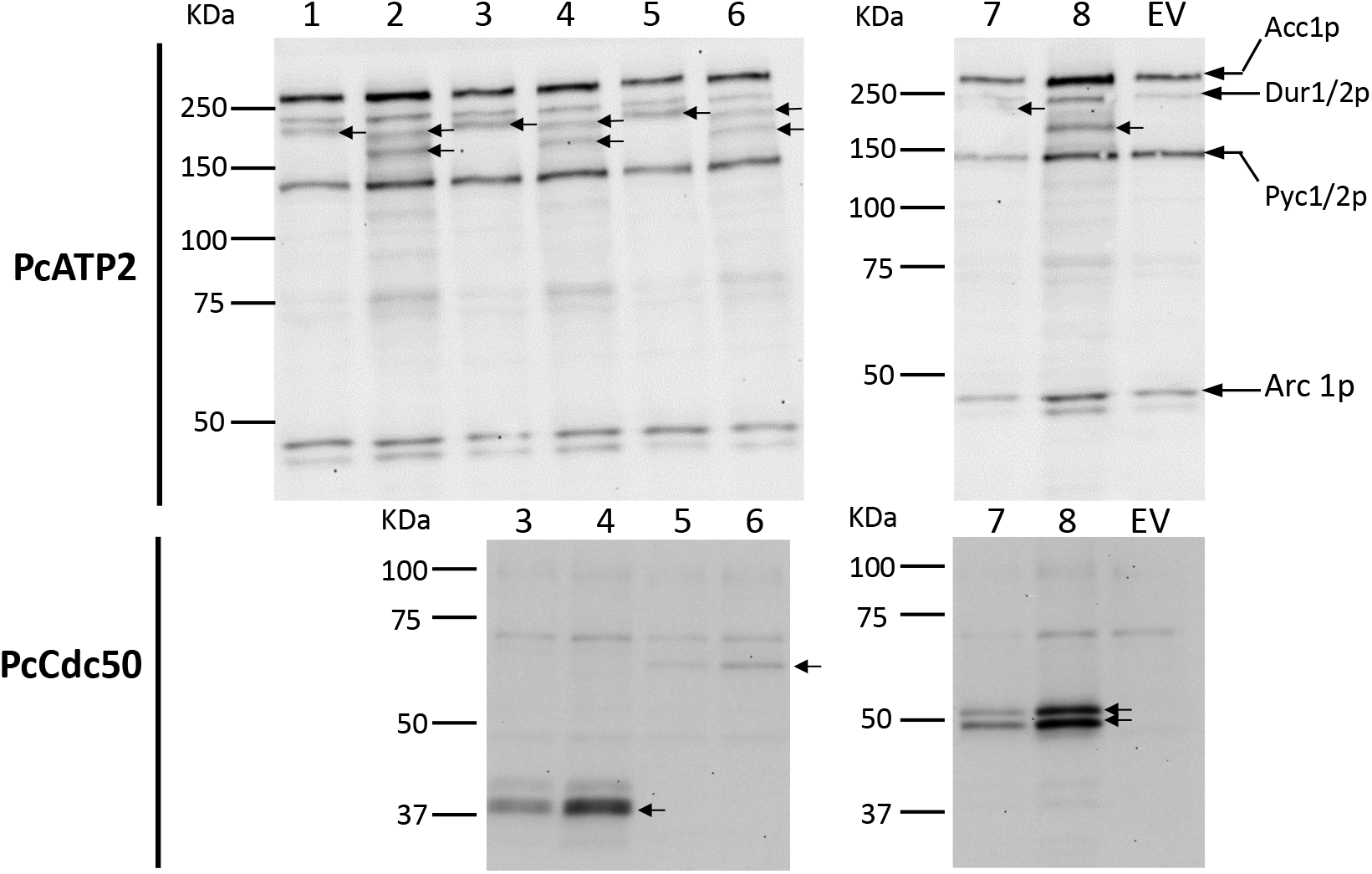
Analysis of PcATP2 expression in *S. cerevisiae* membranes, alone or co-expressed with each of the three putative PcCdc50 subunits. 5 μg of total protein were loaded on each line. Top panels, western blot revealed with the probe against BAD. Bottom panels, western blots revealed with the HisProbe™ to detect the 10xHis tag. Gel bands corresponding to PcATP2 and the *P. chabaudi* Cdc50 subunits are indicated by arrows. *EV:* Empty Vector, membranes expressing no proteins (negative control). *Lane 1:* single expression of PcATP2-BAD. *Lane 2:* single expression of BAD-PcATP2. *Lane 3:* co-expression of PcATP2-BAD and PcCdc50B-His. *Lane 4:* co-expression of BAD-PcATP2 and PcCdc50B-His. *Lane 5: c*oexpression of PcATP2-BAD and PcCdc50C-His. *Lane 6:* co-expression of BAD-PcATP2 and PcCdc50C-His. *Lane 7*: co-expression of PcATP2-BAD and PcCdc50A-His. *Lane 8:* co-expression of BAD-PcATP2 and PcCdc50A-His. The theoretical molecular weight mass of PcATP2-BAD or BAD-PcATP2 is 180 kDa. The theoretical molecular weight masses of PcCdc50B-His, PcCdc50C-His and PcCdc50A-His are, respectively, 45, 61 and 51 kDa. Biotinylated *S. cerevisiae* proteins are indicated in the EV lane: Acc1p, acetyl-CoA carboxylase; Dur 1/2p, urea carboxylase; Pyc 1/2p, pyruvate carboxylase isoforms 1 and 2, and Arc 1p, complex acyl-RNA.

We also examined the expression yield in two different membrane fractions obtained from differential centrifugation of BAD-PcATP2 co-expressing with either, PcCdc50A-His or PcCdc50B-His. We detected the presence of PcATP2 and the two PcCdc50 subunits in the high-density or P2 fraction, and in the low-density or P3 fraction (Figure 2). Notably, PcATP2 changed its relative distribution between P2 and P3 depending on the co-expressed PcCdc50 subunit (Figure 2). That is, while the relative amount of PcATP2 in P2 or P3 was similar when co-expressed with PcCdc50B (Figure 2, left pannels), PcATP2 seems to be mostly accumulated in P2 membranes when co-expressed with PcCdc50A (Figure 2, right panels). In addition, the relative distribution of PcATP2 between P2 and P3 membranes followed the same relative distribution as its co-expressed PcCdc50 protein (Figure 2, bottom panels).

**Figure 2.**
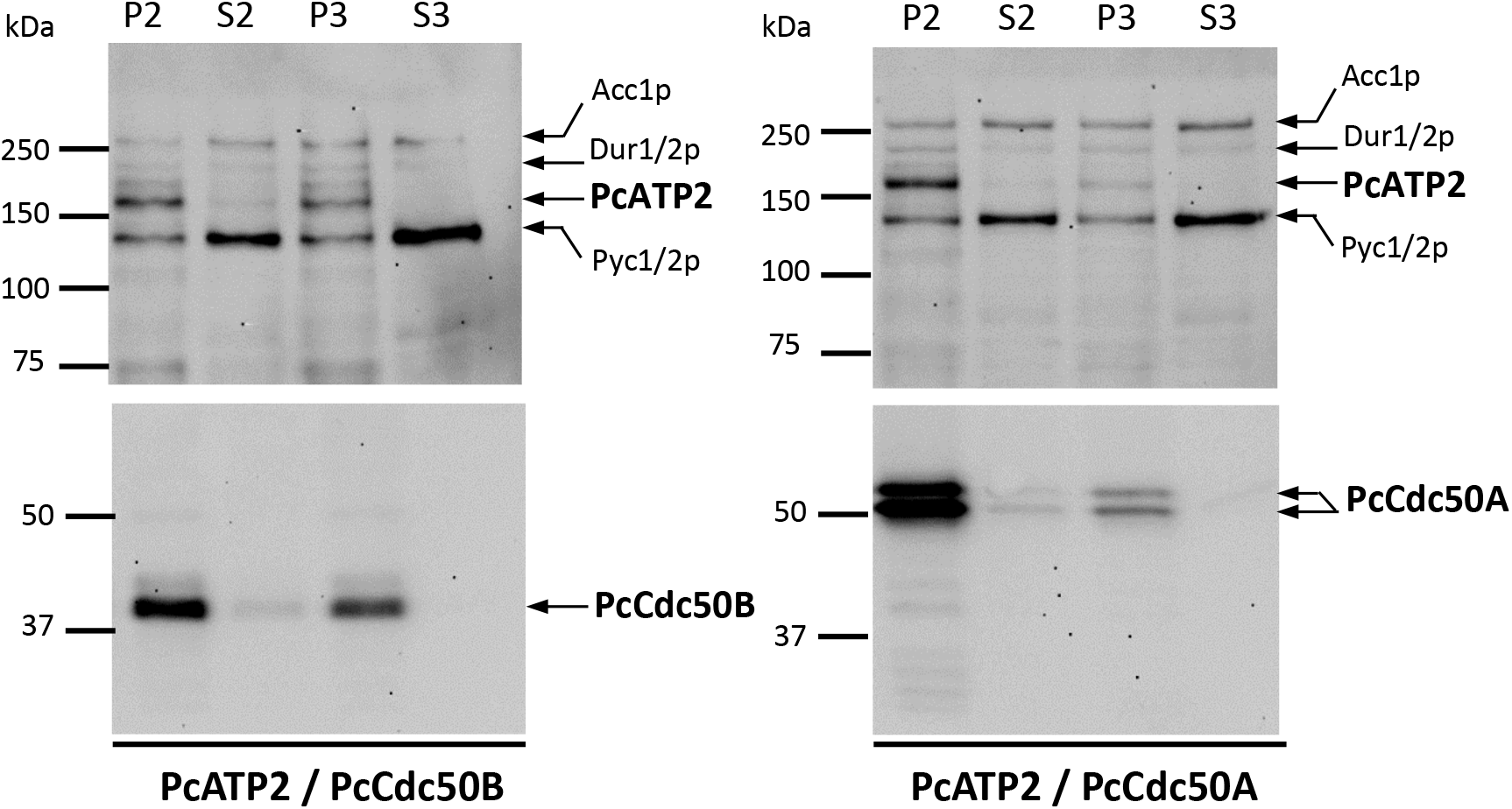
Analysis of membrane fractions of *S. cerevisiae* co-expressing PcATP2 with either PcCdc50B or PcCdc50A. Top panels, western blots revealed with the probe against the BAD. Bottom panels, western blots revealed with the HisProbe™ to detect the 10xHis tag. Left panels, membrane fraction co-expressing BAD-PcATP2 and PcCdc50B-His. Right panels, membrane fraction co-expressing BAD-PcATP2 and PcCdc50A-His. The bands corresponding to BAD-PcATP2, PcCdc50B-His and PcCdc50A-His are indicated. P2 and S2 are, respectively, the membrane pellet and the supernatant obtained after centrifugation at 20000 x g. P3 and S3 are, respectively, the membrane pellet and the supernatant obtained after centrifugation at 125000 x g. Each line was loaded with the corresponding membrane fraction or supernatant containing 1 μg of total protein. Biotinylated *S. cerevisiae* proteins are also indicated: Acc1p: acetyl-CoA carboxylase, Dur 1/2p: urea carboxylase, and Pyc 1/2p: pyruvate carboxylase isoforms 1 and 2.

We screened a battery of eleven detergents to solubilize PcATP2 and the co-expressed Cdc50 proteins from both, P2 and P3 membranes. PcATP2 and Cdc50 proteins in P2 membranes could not be solubilized under any of the tested experimental conditions, with the exception of the rather denaturing n-dodecylphosphocholine (Fos-Choline-12) (Chipot et al. 2018), strongly suggesting a folding defect of the P2-containing proteins (Thomas and Tate 2014). In contrast, a fraction of PcATP2 and PcCdc50B from P3 membranes was solubilized in N-dodecyl-β-D-maltopyranoside (DDM) and, with less efficiency, in Lauryl maltose neopentyl-glycol (LMNG) or Octaethylene glycol monodecyl ether (C12E8) (Figure supplement 3). Importantly, the solubilization efficiency of these detergents was highly improved by the presence of the cholesterol derivative, cholesteryl hemisuccinate (CHS) (Figure supplement 3). Due to the low amount of BAD-PcATP2 co-expressed with PcCdc50A in P3 (Figure 2), the immunodetection of the DDM/CHS solubilized fraction of BAD-PcATP2 from P3 membranes was difficult (Figure supplement 3, panel E). Interestingly, we found that the lower band of PcCdc50A (indentified in the next section as the non-glucosylated form of PcCdc50A, Figure 3) was the only one co-solubilized with PcATP2 (Figure supplement 3, panel F). This suggests that glycosylation of PcCdc50A might affect the folding and/or membrane trafficking during biogenesis, explaing (at least, in part) the preferential localization of PcCdc50A together with its co-expressed PcATP2 partner in P2 membranes (Figure 2, right panels).

**Figure 3.**
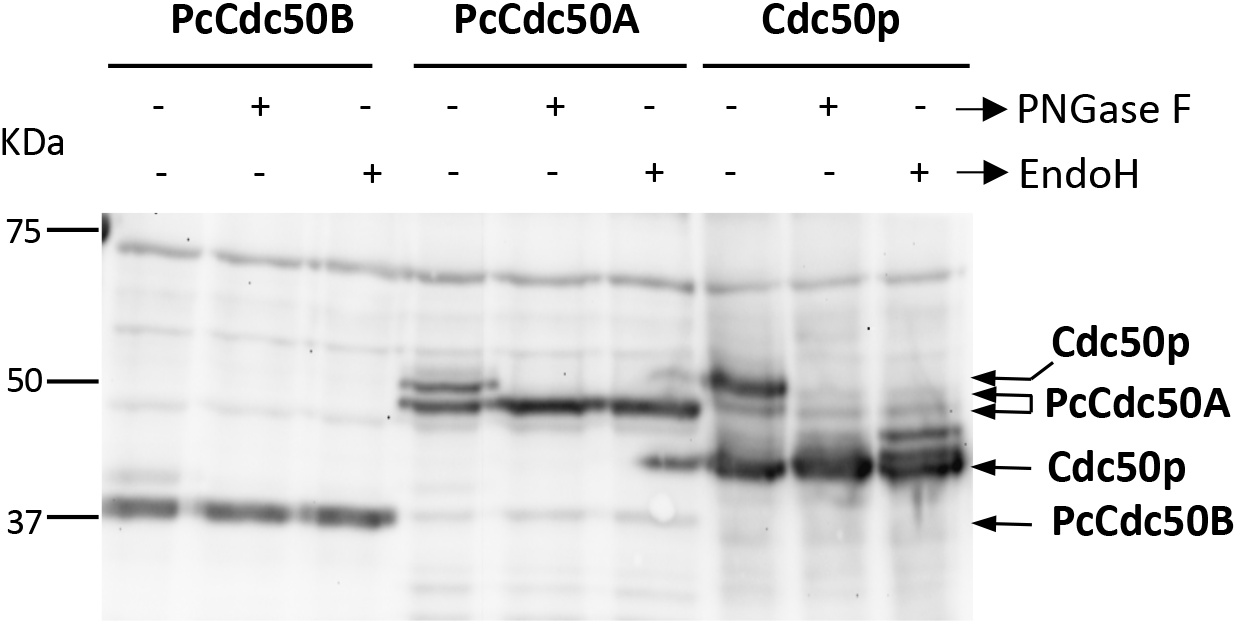
Deglycosylation assays of PcCdc50B and PcCdc50A co-expressed with PcATP2. Membranes co-expressing PcATP2/PcCdc50B, PcATP2/PcCdc50A or Drs2p/Cdc50p were enzymatically deglycosylated with PNGase F or EndoH, as indicated. 10 μg total protein were loaded in the experiments with PcATP2/PcCdc50B and PcATP2/PcCdc50A, whereas 5 μg of total protein were loaded in the control experiment with Drs2p/Cdc50p. The western blots were revealed with the HisProbe™ to detect the 10xHis tag.

### 3. N-Glycosylation of PcCdc50A and PcCdc50B expressed in *Sacharomyces cerevisiae*

As seen in Figures 1 and 2, PcCdc50A-His presented two electrophoretic bands, suggesting glycosylation events typical of Cdc50 proteins. Glycosylation of Cdc50 proteins plays often important roles in the stability and the activity of the P4-ATPase/Cdc50 complex (Costa et al. 2016; García-Sánchez et al. 2014). Using the online server NetNGlyc (http://www.cbs.dtu.dk/services/NetNGlyc/), one potential glycosylation site was predicted at position N260 within the FNDT motif of PcCdc50A. This motif is conserved in Cdc50 proteins from others organisms (García-Sánchez et al. 2014; Poulsen et al. 2008), including the *S. cerevisiae* Cdc50 protein, Cdc50p, whose glycosylation state has already been observed when overexpressed in *S. cerevisiae* (Jacquot et al. 2012). In contrast, no potential N-glycosylation sites were predicted in the sequence of PcCdc50B. To analyze the possible N-glycosylation of PcCdc50A and PcCdc50B when expressed in *S. cerevisiae*, we performed enzymatic deglycosylation assays in membranes co-expressing BAD-PcATP2/PcCdc50B-His or BAD-PcATP2/PcCdc50A-His, using two deglycosydases: the amidase PNGase F and the N-glycosydase EndoH. As control, we used Cdc50p co-expressed with its *S. cerevisiae* P4-ATPase partner, Drs2p (BAD-Drs2p/Cdc50p-His). Consistent with previous studies (Jacquot et al. 2012), Cdc50p displayed two major bands in the western blot (Figure 3); the fully glycosylated fraction (~50 kDa) and the nonglycosylated one (~37 kDa). After incubation with either PNGase F or EndoH, the glycosylated band of Cdc50p disappeared while the lowest or non-glycosylated band remained intact. With regard the *Plasmodium* proteins, the band corresponding to PcCdc50B remained unaltered after enzymatic treatment (Figure 3). Conversely, the upper band of PcCdc50A disappeared after incubation with either PNGase or EndoH, while the lowest band became more intense with the same electropohoretic mobility (Figure 3), thus confirming the N-glysosylation of PcCdc50A predicted from the amino acid sequence. Of note, in the western blot of PcCdc50B there is an extra faint band just above the main band of PcCdc50B untreated protein, which disappeared after enzymatic digestion (Figure 3). Critically, we cannot discard a possible O-glycosilation form of PcCdc50B since potential O-glycosylation sites are predicted at positions 153, 246 and 247 of PcCdc50B, although these sites are not conserved in other *Plasmodium* Cdc50B orthologs. In any case, this band seems to represents a very low population of PcCdc50B, and it is almost absent after detergent solubilization (Figure supplement 3, panel B).

### 4. PcATP2 associates with PcCdc50B and PcCdc50A in detergent micelles

To assess the association of PcATP2 with PcCdc50B or PcCdc50A even after detergent solubilization, we performed immunoprecipitation studies. We fused the superfolder green fluorescent protein (GFP) (Pédelacq et al. 2006) at the C-terminal end of PcATP2 followed by the BAD (PcATP2-GFP-BAD). We chose to tag PcATP2 at the C-terminal end rather than the N-terminal like the previous experiments because the GFP is a more sensitive folding reporter when fused at the C-terminal end of the target protein (Rodríguez-Banqueri et al. 2016). The presence of both co-expressed proteins was confirmed by western blot (Figure supplement 4). P3 membranes co-expressing PcATP2-GFP-BAD/PcCdc50B-His or PcATP2-GFP-BAD/PcCdc50A-His were solubilized with DDM/CHS, and detergent-solubilized PcATP2-GFP-BAD was trapped in agarose beads coupled to an anti-GFP nanobody (or GFP-Trap^®^) (Rothbauer et al. 2008). After washing the beads, bound proteins were eluted with a low-pH buffer and analyzed by western blot. As expected, PcATP2-GFP-BAD was detected in both elution fractions (Figure 4, lane 4 top panels). Moreover, PcCdc50B-His and PcCdc50A-His also coeluted with PcATP2-GFP-BAD in their respective experiments (Figure 4, lane 4 bottom panels), demonstrating that PcATP2 interacts with either, PcCdc50B or PcCdc50A in detergent micelles. The western blot signals of eluted PcATP2-GFP and PcCdc50A-His were much weaker than the ones of the PcATP2-GFP/PcCdc50B-His experiment, in concordance with their relative amounts in the P3 membrane fraction (Figure 2 and Figure supplement 4). As control, we performed the same experiment using P3 membranes co-expressing a non GFP-tagged PcATP2 with either, PcCdc50B (PcATP2-BAD/PcCdc50B-His) or PcCdc50A (PcATP2-BAD/PcCdc50A-His). In this case, none of the PcCdc50 proteins were detected in the elution fraction (Figure 4, lane 8 bottom panels), thus discarding a false positive result due to a non-specific binding of PcCdc50B-His or PcCdc50A-His to the GFP-Trap^®^ agarose beads.

**Figure 4.**
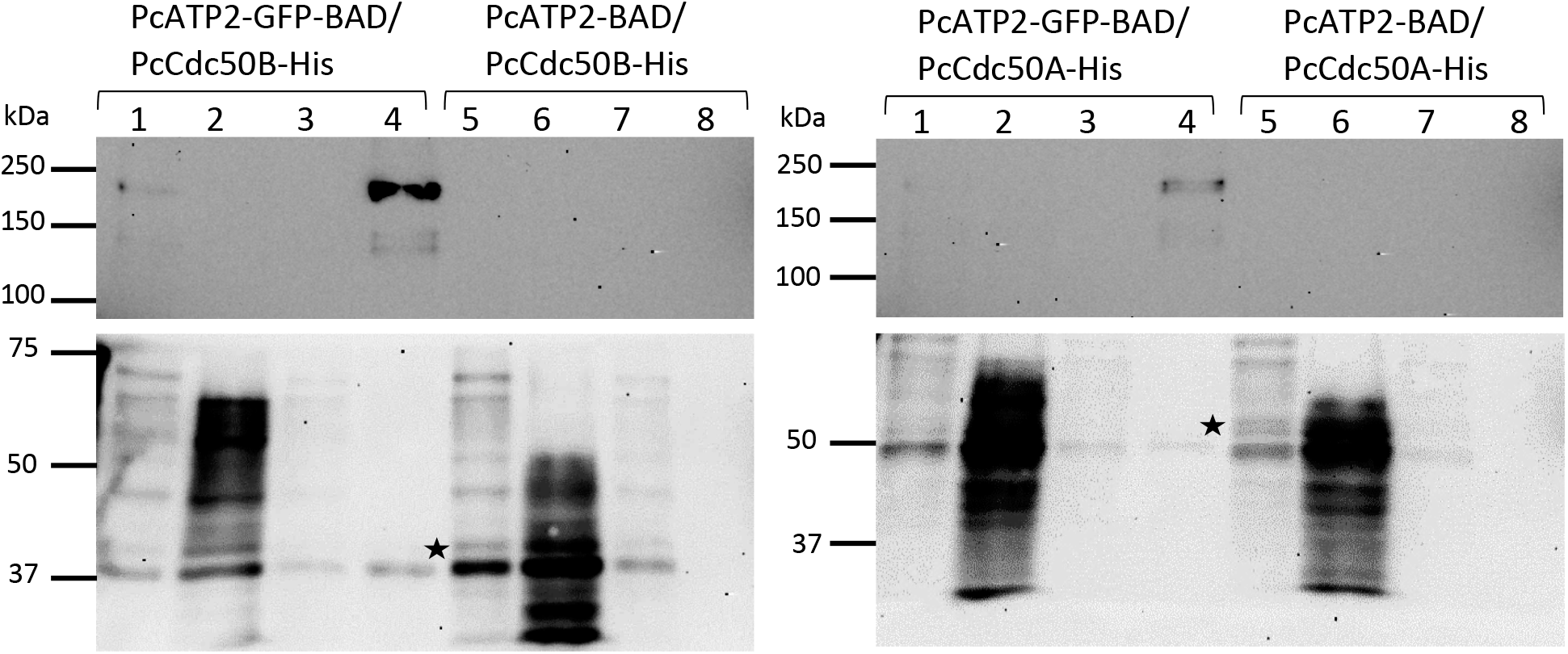
Co-immunoprecipitation of PcATP2-GFP-BAD with PcCdc50B-His or PcCdc50A-His. Lanes *1* to *4* correspond to the different steps of the assay with membranes co-expressing PcATP2-GFP-BAD and either, PcCdc50B-His or PcCdc50A-His: (*1*) input, detergent-solubilized membranes, (*2*) flow-through or non-bound protein fraction, (*3*) washings, and (*4*) elution. Lanes *5* to *8* are the equivalent lanes *1* to *4* with membranes co-expressing PcATP2-BAD and either, PcCdc50B-His or PcCdc50A-His. The blots on the top panel were revealed with an antibody against the GFP to detect PcATP2-GFP-BAD, and the bottom one with the HisProbe™ to detect PcCdc50B-His and PcCdc50A-His (labelled with an asterisk).

We also analyzed PcATP2 and PcCdc50B association by fluorescent size-exclusion chromatography (FSEC). We used the GFP fluorescence to monitor the elution profile of DDM/CHS-solubilized PcATP2 in a gel-filtration column, and analyzed the degree of monodispersity (or stability) in detergent micelles of PcATP2 in the presence or in the absence of PcCdc50B. Detergent-solubilized PcATP2-GFP-BAD eluted as a nearly symmetric peak centered at ~11.5 ml (Figure 5A). The second peak at ~16 ml elution corresponded to the light-scattering of empty DDM/CHS micelles due to the high instrumental gain used in these samples (see shadow chromatogram in Figure 5A that corresponds to DDM/CHS-solubilized membranes expressing no GFP-tagged protein). The shape and retention time of PcATP2 did not change significatively between single expression and co-expression experiments (Figure 5A), suggesting a reasonably stability of PcATP2 in DDM/CHS micelles both, in the presence and in the absence of PcCdc50B. PcCdc50B-His also co-eluted with PcATP2-GFP-BAD as revealed by the western blot analysis of the eluted fractions along the PcATP2-GFP-BAD elution peak (Figure 5A, upper panel), following a similar elution profile than PcATP2-GFP-BAD (Figure 5A). We also fused the GFP at the C-terminal end of PcCdc50B (PcCdc50B-GFP-His) and performed FSEC analysis of DDM/CHS-solubilized membranes expressing PcCdc50B-GFP-His alone or co-expressed with PcATP2-BAD (devoid of the GFP). The elution profile of PcCdc50B-GFP-His in the absence of PcATP2 was relatively broad and asymmetric (Figure 5B), an indication of poor stability in DDM/CHS. Remarkably, when co-expressed with PcATP2-BAD, the elution profile of PcCdc50B-GFP-His became narrower and more symmetric, with a main elution peack centered at ~14.5 ml (Figure 5B). No peack at ~16 ml corresponding to empty micelles appeared because we used a lower detector gain in the instrumental set up. Our data suggested a stabilization role of PcATP2 on PcCdc50, at least, after solubilization in DDM/CHS detergent micelles. It is unlikely that PcATP2-BAD is also co-eluting with PcCdc50B-GFP-His at this ~14.5 ml peack (Figure 6B) as PcATP2-GFP-BAD alone or co-expressed with PcCdc50B-His eluted at ~11.4 ml (Figure 5A). As the fluorescence intensity at the main elution peak of PcCdc50B-GFP-His (310 mA at 14.5 ml, Figure 5B) was approximately ten times higher than the one of PcATP2-GFP-BAD (37 mA at 11 ml, Figure 5A), we made the assumption that the amount of detergent-solubilized PcCdc50B-GFP-His was approximately ten times higher than the amount of PcATP2-GFP-BAD when both proteins were co-expressed. In this scenario, and assuming the 1:1 (mol/mol) stoichiometry of the PcATP2/PcCdc50B complex typical of P4-ATPase/Cdc50 heterodimers, in the normalized FSEC presented in Figure 5B we would expect a maximum fluorescent intensity corresponding to the elution of PcATP2-BAD/PcCdc50B-GFP-His complex of 0.1 at ~11.4 ml, which would be hidden by the shoulder of PcCdc50B-GFP-His elution. In summary, FSEC experiments confirmed the association of PcATP2 with PcCdc50B, as well as the stability of this complex in DDM/CHS micelles as judged by the FSEC’s elution profile.

**Figure 5.**
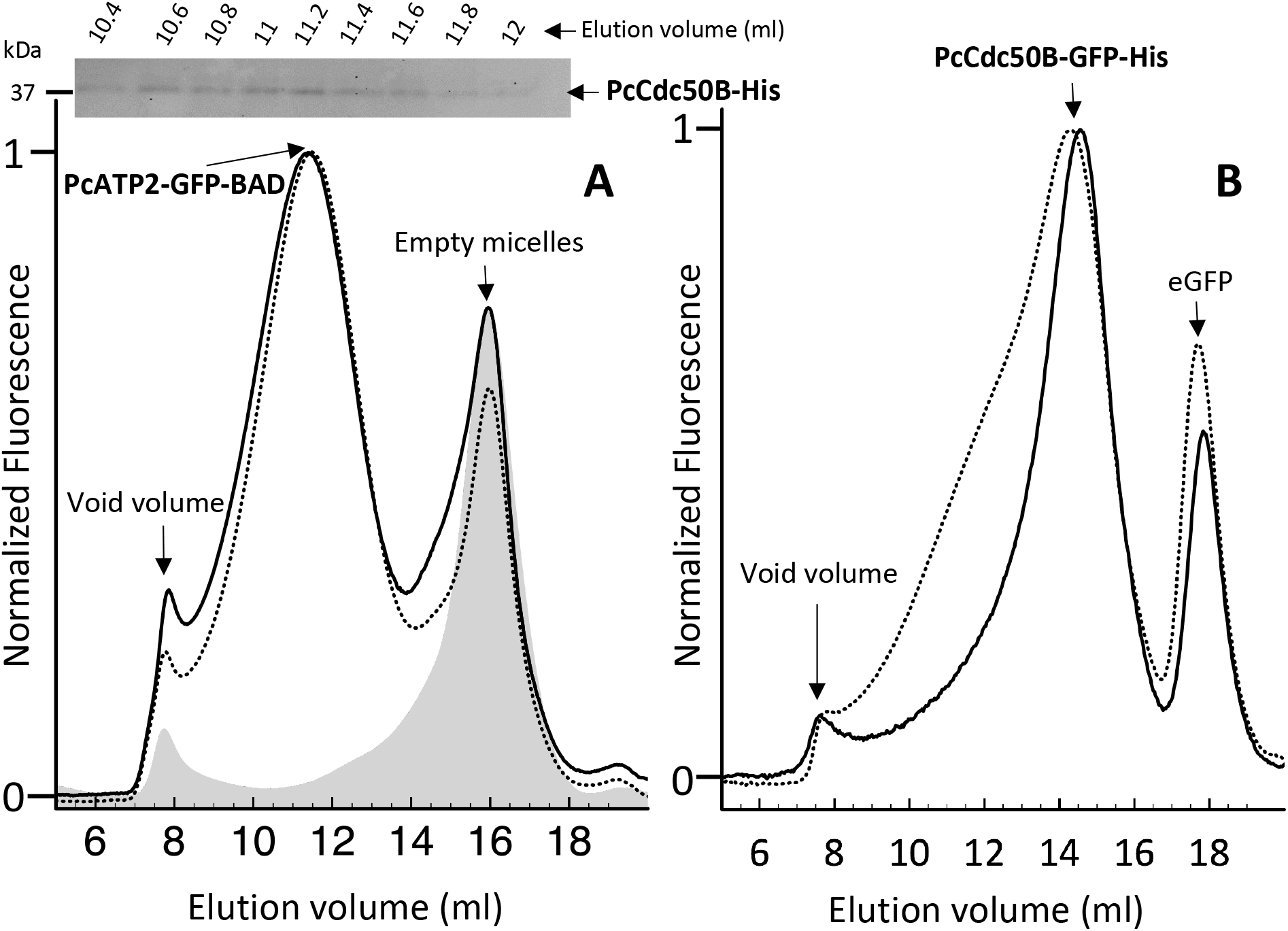
Fluorescence-detection Size Exclusion Chromatography of detergent solubilized PcATP2 and PcCdc50B. Membranes were solubilized with 1 % (w/v) DDM, 0.2 % (w/v) of CHS, and the supernatant after ultracentrifugation was loaded into a Superose 6 10/300 GL gel-filtration column equilibrated with 20 mM Tris-HCl pH 7.8, 150 mM NaCl, 10 % (v/v) Glycerol, 0.1 mg/mL DDM, 0.02 mg/mL CHS, and connected to a fluorescence detector. Panel *A*: Normalized FSEC profiles of single-expression of PcATP2-GFP-BAD (dotted lines) and co-expression of PcATP2-GFP-BAD with PcCdc50B-His (solid line). In grey shadow, profile of membranes expressing no protein (control). Inside panel *A*, analysis of PcCd50B-His co-elution with PcATP2-GFP-BAD. Collected fractions of 200 μl between 10.4 to 12 ml of the PcATP2-GFP-BAD elution were analyzed by western blot using the HisProbe™ to detect the presence of PcCdc50B-His. Panel *B*, Normalized FSEC profiles of single-expression PcCdc50B-GFP-His (dotted line) and co-expression PcATP2-BAD/PcCdc50B-GFP-His (solid line).

**Figure 6.**
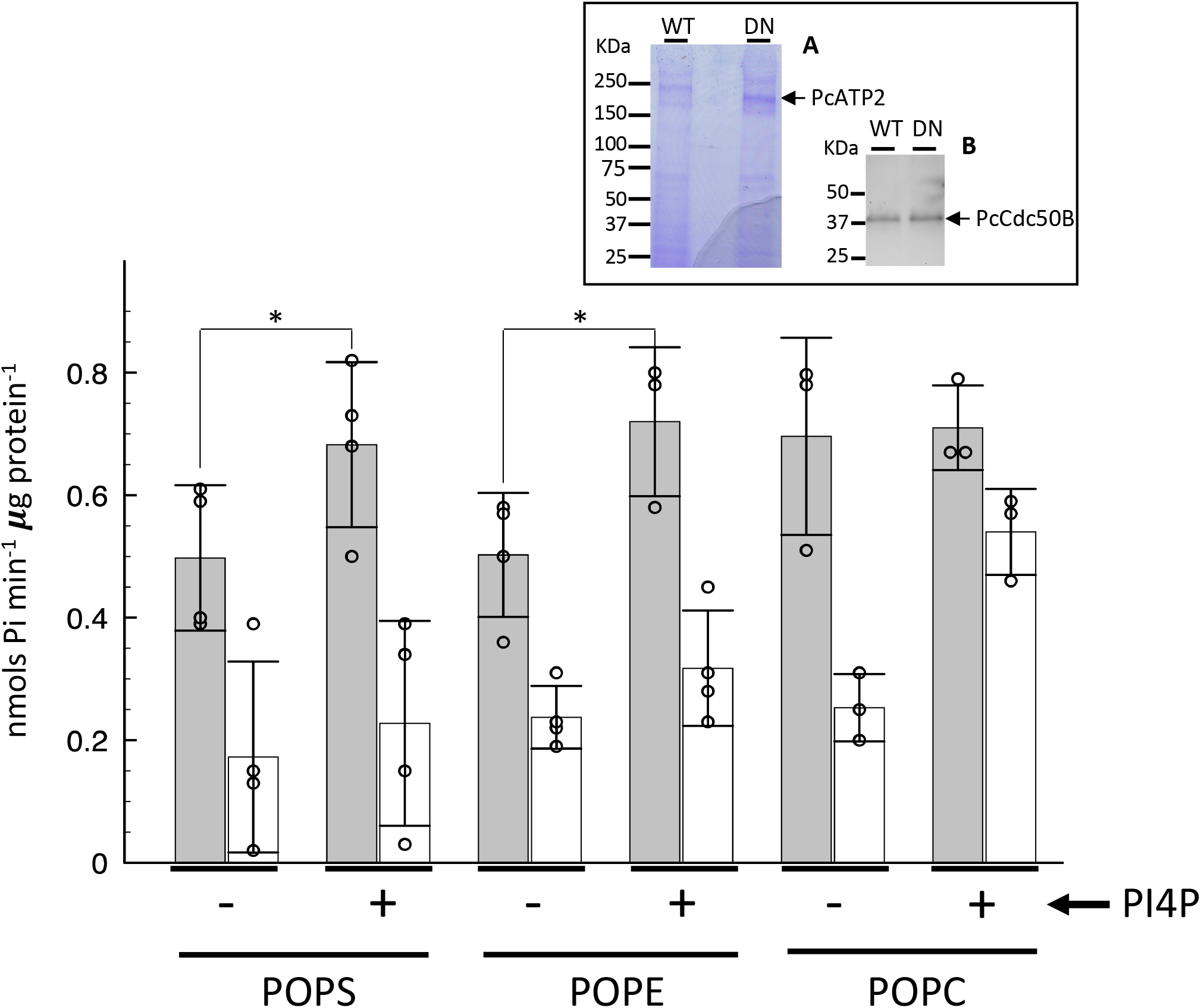
Lipid-dependent ATPase activity of PcATP2/PcCdc50B complex. The apparent ATPase activity of the PcATP2/PcCdc50B complex was measured by spectrophotometric quantification of the released inorganic phosphate in the presence of the indicated lipids. POPS, 1-palmitoyl-2-oleoyl-sn-glycero-3-phospho-L-serine, POPE, 1-Palmitoyl-2-oleoyl-sn-glycero-3-phosphoethanolamine, POPC, 1-palmitoyl-2-oleoyl-sn-glycero-3-phosphocholine. As indicted, ATPase assays were also performed in the presence (+) or in the absence (-) of phosphatidylinositol-4-phosphate (PI4P). Dark bars and white bars represent, respectively, the activity of the wild-type and the functionally-impaired D596N mutant at each experimental condition. The reaction was performed at 37°C during 25 minutes, and initiated by addition of 1 mM ATP-Mg^2+^. The bars display the average value of n=3 or n=4 experimental datapoints, and the indicated error is the standard deviation. The upper panel displays the Coomassie-stained (*A*) or the western blot using the HisProbe™ to detect the 10xHis tag (*B*) SDS-PAGE of the acid-eluted content of the immobilized wild-type (WT) or D596N mutant (DN) agarose beads used for this experiments. * P<0.05 (Paired T-test). Note that the DN mutant of PcATP2 runs slightly faster because it lacks the BAD domain.

### 5. Detergent-solubilized PcATP2/PcCdc50B catalyzes ATPase activity in the presence of POPS, POPE and POPC, and senses PI4P

In detergent micelles, P4-ATPases hydrolyze ATP only in the presence of their phospholipid substrates (Coleman, Kwok, and Molday 2009; Zhou, Sebastian, and Graham 2013; Theorin et al. 2019). This property not only permits the partial evaluation of the activity of the transporter along several steps of the transport cycle, but also it allows to assess substrate specificity. We assayed the functional properties of recombinant PcATP2-GFP-BAD/PcCdc50B-His complex by measuring the lipid-stimulated capacity of PcATP2 to catalyze ATP hydrolysis. Detergent-solubilized PcATP2-GFP-BAD/PcCdc50B-His was immobilized in home-made GFP-Trap^®^ beads, and the liberated inorganic phosphate upon ATP hydrolysis was quantified spectrophotometrically after 25 min at 37°C in the presence of three different phospholipids. To reduce the potential ATPase activity from traces of the endogenous F0F1 ATPase, we added 1 mM of NaN3, a potent inhibitor of this enzyme (Centeno et al. 1994). In addition, as negative control, we generated a predicted non-functional mutant of PcATP2 after replacing the aspartate residue at position 596 by asparagine (Figure supplement 1). This aspartate is part of the conserved P-type ATPase motif DKTG, and undergoes phosphorylated and de-phosphorylated during each transport cycle (Palmgren and Nissen 2011). Therefore, the resulting PcATP2-D596N-GFP mutant is expected to be fully deficient in ATP hydrolysis. PcATP2-GFP wild-type and PcATP2-D596N-GFP expressed at similar yields (Figure supplement 4) and the presence of both PcATP2 variants and the co-expressed PcCdc50B-His in the home-made GFP-Trap^®^ beads were confirmed by SDS-PAGE (Figure 6, inserted panel). Although both PcATP2 versions were visible by Coomassie staining (Figure 8, inserted panel A), PcCdc50B-His was only visible after immunoblotting (Figure 8, inserted panel B). Inmobilized PcATP2-GFP-BAD/PcCdc50B-His in GFP-Trap^®^ beads catalyzed ATPase activity in the presence of the phospholipids, POPS, POPE and POPC (Figure 6), with an apparent ATPase activity between ~0.5 and 0.7 nmols Pi min^−1^ μg protein^−1^. In the same conditions, the activity of the functionally-impaired mutant, PcATP2-D596N-GFP/PcCdc50B-His ranged from ~0.2 to 0.4 nmols Pi min^−1^ μg protein^−1^ (Figure 6), therefore the heterologously produced PcATP2 catalizes ATPase activity. Fluctuations of the ATPase activity between samples are normally observed in other detergent-solubilized P4-ATPases (Theorin et al. 2019), likely due to the slightly different content of cellular lipids carried by the detergent-solubilized proteins between different experiments. The C-terminal end of PcATP2 and its *Plasmodium* orthologs contains the equivalent residues Tyr1235 and His1236 of Drs2p, recently identified as a phosphatidylinositol-4-phosphate PI4P recognition motif (Timcenko et al. 2019) (Figure supplement 5). In Drs2p, PI4P binding induces an increase of the substrate-stimulated ATPase activity (Jacquot et al. 2012). Similarly, when PI4P was present in addition to either POPS or POPE, we also observed an increase of the apparent ATPase activity of the PcATP2-GFP-BAD/PcCdc50B-His complex, while the PI4P effect in the PcATP2-D596N-GFP-BAD mutant was much more reduced or absent (Figure 6). Unfortunately, in our experimental set up we could not raise any conclusion with regard to a specific increase of POPC-induced PcATP2 activity upon PI4P addition, as the apparent activity in the presence of PI4P also increased in the sample containing the PcATP2-D596N-GFP-BAD mutant (Figure 6). In summary, the detergent-solubilized PcATP2/PcCdc50 complex produced in yeast is functionally competent as its catalizes ATPase activity in the presence of POPS, POPE and POPC, as well as it is upregulated by PI4P in the presence of POPS and POPE.

## DISCUSSION

The essential and irreplaceable putative lipid flippase activity of ATP2 (Kenthirapalan et al. 2016; Bushell et al. 2017; Zhang et al. 2018) confirms that lipid homeostasis is paramount for the parasite’s development, representing new opportunities for therapeutic intervention. The challenge now is to understand why and when the transport activity of ATP2 and the other putative P4-ATPases are required within the complex life-cycle of the malaria parasite. The intracellular development inside the parasitized cell depends on an initial heavy supply of lipids to build the different membrane compartments, together with efficient mechanisms to ensure a suitable lipid composition on each membrane, as ~75 % of all lipids exhibit significant variations on its relative amount along parasite’s development, particularly during gametogenesis (Gulati et al. 2015; Tran et al. 2016). The *Plasmodium* parasite has a limited capacity of synthesizing fatty acids and phospholipids, and therefore, many lipids are scavenged from the host membrane or from the serum (Kilian et al. 2018) where P4-ATPases might have a relevant role.

The production of recombinant malaria transporters has been pivotal in malaria research, allowing the identification of substrates and inhibitors (Juge et al. 2015; Ferreira et al. 2011; David-Bosne et al. 2013), providing protein sample for structural studies (Qureshi et al. 2020; Kim et al. 2019) or for discarding the role of PfATP6 as artemisinin drug-target (Cardi et al. 2010; Arnou et al. 2011; David-Bosne et al. 2016). To unravel the structural and functional features of ATP2 we screened three ATP2 orthologs for *S. cerevisiae* expression. In the present study we only succeeded to express the *P. chabaudi* ATP2, PcATP2. This expression system was found to be successful in our group for PfATP6 (Cardi et al. 2010), a yeast P4-ATPase (Azouaoui et al. 2014), and a rabbit Ca^2+^ P-type ATPase, which could be crystalized after purification (Jidenko et al. 2005). P-type ATPases encoded by apicomplexan parasites contain non-conserved amino acid stretches within the two large cytoplasmic regions connecting TMDs 1 and 2 and TMDs 4 and 5 (Figure 7), enclosing a high density of positively-charged and asparagine residues, a potential handicap for heterologous expression (Fujita et al. 2010). These extensions are absent in many other P-type ATPases, and as judged by a 3D model of PcATP2 based on the recent cryo-EM structure of Drs2p and ATP8A1 (Hiraizumi et al. 2019; Timcenko et al. 2019), they contain non-structured regions (Figure 7). Nevertheless, PcATP2 is a fair paradigm for the biochemical characterization of *Plasmodium* ATP2 due to its high conservation in *Plasmodium* species (Figure supplement 1), Co-immunoprecipitation studies (Figure 4), together with membrane fraction co-localization (Figure 2) and detergent-solubilization screenings (Figure supplement 3, panels *E* and *F*) demonstrated that the yeast-produced PcATP2 is able to associate with both PcCdc50B and PcCdc50A, two of the three *Plasmodium-encoded* Cdc50 isoforms. This promiscuous capacity to associate with more than one Cdc50 β-subunit has been observed in other P4-ATPases (Paulusma et al. 2008; Van Der Velden et al. 2010), although the physiological meaning of this dual interaction remains to be determined. We also found that *S. cerevisiae-expressed* PcCdc50A was N-glycosylated (Figure 3). In contrast, PcCdc50B presented no N-glycosylated when expressed in *S. cerevisiae* (Figure 3), although possible O-glycosylation could not be rouled out (Figure 3 and Figure supplement 2). As N-glycosylation is a typical signature of Cdc50 proteins (Costa et al. 2016), PcCdc50B would be, to the best of our knowledge, the first characterized Cdc50 protein containing no N-glycosylations. The role of N-glycosylations of Cdc50 proteins seems to be organism-dependent. While in the *Leishmainia* parasites, Cdc50 glycosylation affects the catalytic activity of the P4-ATPase/Cdc50 complex (García-Sánchez et al. 2014), N-glycosylation of the Cdc50 β-subunit of the human ATP8A2 is required to form a stable complex with the P4-ATPases, with no apparent role on its catalytic activity (Coleman and Molday 2011). In *Plasmodium*, with the exception of the largely glycosylated proteins exported at the plasma membrane of the infected erythrocyte, glycosylation of proteins is rare, despite the fact that the parasite encodes enzymes for N- and O-glycosylation, and predicted glycosylation sites are present in numerous *Plasmodium-encoded* proteins (Cova et al. 2015). Interestingly, the *P. yoeli* Cdc50A expressed in both gametocytes and ookinetes does not seem to be glycosylated as judged by the western blots bands ((Gao et al. 2018), and Jing Yuan, personal communication). Clearly, due to its potential functional and/or stabilizing role, further studies to unravel the glycosylation state of Cdc50A in the parasite are necessary.

**Figure 7.**
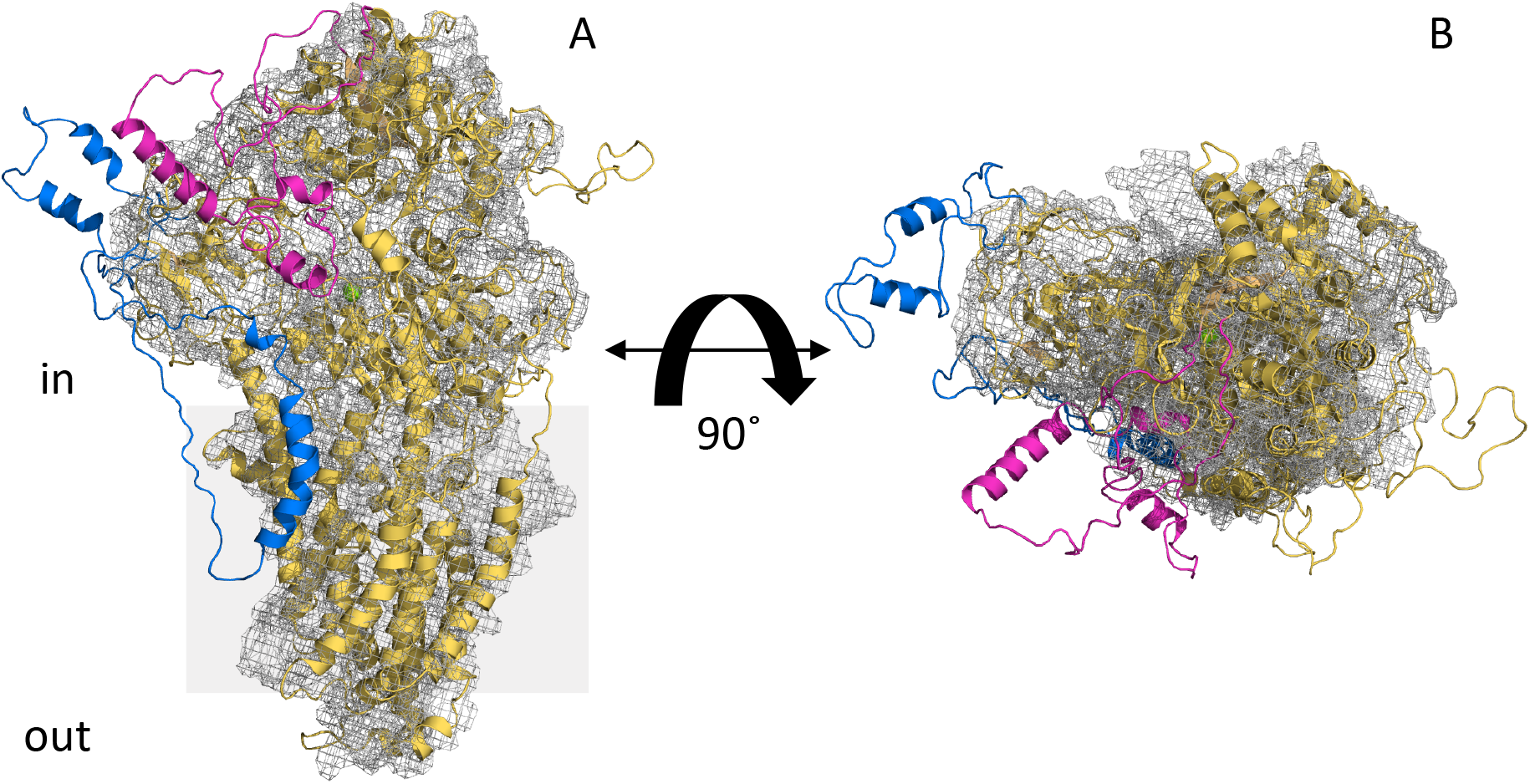
Alignment of the cryo-EM structure of Drs2p and a 3D structural model of PcATP2. The 3D structural model of PcATP2 (cartoon representation) was aligned with the cryo-EM structure of the autoinhibited conformation of Drs2p (PDB ID 6roh, mesh representation). Protein domains of PcATP2 in the cytoplasmic region between TMDs 2 and 3 that are absent in Drs2p are depicted in blue. Protein domains of PcATP2 in the cytoplasmic region between TMDs 4 and 5 that are absent in Drs2p are depicted in magenta. (*A*) Lateral perspective of the molecule. (*B*) Cytoplasmic view after 90° rotation of *A* through the x axis. The grey shadow square in *A* represents the putative position of the surrounding membrane.

FSEC experiments confirmed the assoaciation and reasonable stability of PcATP2 with PcCdc50B in DDM/CHS detergent micelles (Figure 5A). Our data also suggested that the cholesterol analog CHS is important to stabilize this complex since its presence highly improved the yield of co-solubilization of PcATP2 and PcCdc50B with DDM (Figure supplement 3). In the recent cryo-EM structure of the human ATP8A1 in complex with Cdc50a, a CHS molecule was found at the heterodimer interface between TMDs 7 and 10 of ATP8A1 and TMD2 of Cdc50a, suggesting a potential role of CHS on preserving the heterodimer (Hiraizumi et al. 2019). Although *Plasmodium* parasites cannot synthetize *de novo* cholesterol, lipidomic studies have revealed the presence of this lipid in intracellular membrane compartments of infected erythrocytes (Botté et al. 2013), most likely obtained from the host cell-membrane. FSEC experiments also revealed a chaperon or stabilizing role of PcATP2 over PcCdc50B (Figure 5B), in line with a previous observation where the P4-ATPase is required for exiting its Cdc50 partner from the ER after forming the functional P4-ATPases/Cdc50 complex (Van Der Velden et al. 2010).

The PcATP2/PcCdc50B complex produced in *S. cerevisiae* is functionally active with respect substrate recognition and ATP hydrolysis, as it showed lipid-dependent ATP hydrolysis in DDM/CHS micelles (Figure 6). PcATP2/PcCdc50B hydrolyzes ATP in the presence of the amino-phospholipids POPS and POPE, as well as in the presence of the neutral phospholipid POPC (Figure 6), suggesting the capacity of PcATP2 to recognize all three phospholipids. Two amino acid clusters determine, at least in part, the lipid head-group selectivity of P4-ATPases (Hiraizumi et al. 2019). In PcATP2, these clusters correspond to residues Gln65 and Leu66 in TMD1, and Ala455 and Asn456 in TMD4 (Figure supplement 1). PS-selective P4-ATPases normally contain polar residues at these two clusters (two Gln at the corresponding positions 65 and 66 of PcATP2, and two Asn at the equivalent positions 455 and 456 of PcATP2) that stablish H-bonds with PS. In contrast, PC-selective P4-ATPases usually contain non-polar residues at the same positions (Baldridge and Graham 2013; Hiraizumi et al. 2019; Vestergaard et al. 2014). PcATP2 has one polar residue at each cluster (Figure supplement 1) that could be sufficient to coordinate the three different head groups. Although in the ATPase assays it is possible the presence of a small non-functional fraction of PcATP2, it is also possible that PcATP2 is partially autoinhibited using a mechanism similar to its yeast homolog Drs2p (Azouaoui et al. 2017; Timcenko et al. 2019). In fact, the GFAFS motif at the C-terminus of Drs2p involved in enzyme’s autoinhibition by interacting with the nucleotide binding site (Timcenko et al. 2019), is also well conserved in PcATP2 (as well in the other *Plasmodium* ATP2 orthologs), although the length of the C-terminal of the *Plasmodium* sequences is ~60 amino acid shorter (Supplementary Figure 5). In addition, PcATP2 shares with Drs2p some residues in the nucleotide binding domain that directly interact with the GFAFS motif to stabilize the autoinhibitory conformation (Timcenko et al. 2019) (Figure supplement 5). Moreover, and as suggested by the presence of a PI4P binding motif on its amino acid sequence (Figure supplement 5)(Timcenko et al. 2019), PcATP2 is also upregulated by this lipid as we observed a statistically significant increase of POPS and POPE-induced ATPase activity in the presence of PI4P (Figure 6). As for all eukaryotic cells (Y. J. Wang et al. 2003), PI4P is also essential for the parasite since the inhibition of PI4P synthesis in the Golgi apparatus completely blocks the late steps of the intraerythrocytic cycle of *P. falciparum* by interfering with membrane biogenesis around the developing merozoites (McNamara et al. 2013). PI4P is found at the plasma membrane and at the Golgi apparatus in all stages of the erythrocytic cycle (Ebrahimzadeh, Mukherjee, and Richard 2018). Interestingly, the *P. berghei* ATP2 was also localized at the interface parasite/host (Kenthirapalan et al. 2016). It will be relevant to investigate if the lethal effects as a result of depleting PI4P or suppressing ATP2 are somehow connected and linked to the biogenesis of the parasite membranes.

In conclusion, our work describes the first biochemical study of a *Plasmodium* ATP2 in complex with a Cdc50-β subunit, a first step towards the understanding of the essential role of this transporter during malaria infection, and its validation as antimalarial drug target.

## ACKNOWLEGMENTS

We would like to thank Dr Christine Jaxel for all the support and discussions during this project, and to Dr. Thibaud Dieudonné, Dr Guillaume Lenoir, Dr Cédric Montigny, Dr Eugene Diatloff and Dr Jing Yuan for crytically reading the manuscript. We also thank Quentin Rochette for his early work on this project. This work was supported by the ANR grants ANR-14-CE09-0022 to Dr Guillaume Lenoir, and ANR-18-C811-0009-02 to JLVI, the French Infrastructure for Integrated Structural Biology (FRISBI; ANR-10-INSB-05), and the Centre National de la Recherche Scientifique (CNRS). In memory of Prof Ronald H. Kaback.

## MATERIALS AND METHODS

### Molecular cloning

Synthetic cDNAs encoding three putative ATP2 orthologs from *P. falciparum (PF3D7_1219600), P. berghei* (PBANKA_1434800), and *P. chabaudi (PCAHS_1436800)*, and the three Cdc50 subunits encoded per specie: *P. falciparum* (PF3D7_0719500, PF3D7_1133300, and PF3D7_1029400), *P. berghei* (PBANKA_061700, PBANKA_091510, and PBANKA_051340), and *P. chabaudi* (PCHAS_061870, PBANKA_091510, and PBANKA_051340) were ordered to Genscript (Piscataway, NJ). All the cDNAs were codon-optimized by the provider for *S. cerevisiae* expression using the proprietary algorithm OptimumGene™. In all cloning steps, we used the *E. coli* XL1-Blue strain and the Luria-Bertani medium with the appropriate antibiotic.

#### Single expression vectors

The cDNAs encoding the three ATP2 orthologs were cloned in the pYeDP60 expression vector (Azouaoui et al. 2014) between the *EcoRI* and *NotI* restriction sites, yielding the pYeDP60-ATP2-TEV-BAD plasmid that contains in downstream position the tobacco etch virus (TEV) protease cleavage-site coding sequence followed by the biotin acceptor domain (BAD) coding sequence. To obtain the N-terminal BAD tagged ATP2 constructs, pYeDP60-BAD-TEV-ATP2, we amplified by PCR the ATP2 sequences incorporating the *Pmel* and *Sacl* restrictions sites at both, the 5’ and 3’ ends, and cloned into the pYeDP60 vector. The PcATP2-D596N mutant was obtained using the QuikChange II XL site-directed mutagenesis kit (Agilent). The nine Cdc50 subunits sequences were cloned in the pYeDP60 vector between the *EcoRI* and *BamHI* restriction sites. The resulting pYeDP60-Cdc50-TEV-10xHis vectors contain the TEV site followed by a 10xHis tag at the C-terminal end of the proteins.

#### Co-expression vectors

pYedBAD-BAD-Drs2p/Cdc50p-His was cloned previously (Azouaoui et al. 2014). To construct the other co-expression vectors we used a previous strategy (Azouaoui et al. 2016). The fragments containing the cassette Promoter-Cdc50-TEV-His10-Terminator from each pYeDP60-Cdc50-TEV-10xHis vector were amplified by PCR, incorporating the *SbfI* restriction site at both the 5’ and the 3’ ends. The PCR fragments were then cloned at the *SbfI* site of the corresponding species-specific single-expression vectors pYeDP60-ATP2-TEV-BAD and pYeDP60-BAD-TEV-ATP2.

### Introduction of the superfolder Green Fluorescent Protein

The cDNA encoding the superfolder Green Fluorescent Protein (GFP) (Pédelacq et al. 2006) was PCR amplified, introducing a *NotI* site followed by the human Rhinovirus 3C protease sequence (3C-protease) at the 5’, and the *XmaI* site at the 3’. The PCR product was then cloned into the *Notl* and *Xmal* sites of both the single-expression pYeDP60-PcATP2-TEV-BAD and the co-expression pYeDP60-PcATP2-TEV-BAD/PcCdc50B-10xHis vectors to obtain the pYeDP60-PcATP2-3C_Protease-GFP-TEV-BAD and the pYeDP60-PcATP2-3C_Protease-GFP-TEV-BAD/PcCdc50B-10xHis vectors. Using an identical cloning strategy, we also made the same constructs but without the BAD domain by adding a stop codon next to the *XmaI* site, leading to the the pYeDP60-PcATP2-D596N-3C _Protease-GFP/PcCdc50B-10xHis vector. To introduce the sGFP at the C-terminal end of PcCdc50B, we amplified by PCR a DNA fragment encoding the 3C_protease followed by the GFP with *BamHI* sites at both the 5’ and the 3’ ens. The sequence was then cloned at the *BamH1* site of pYeDP60-PcCdc50B-TEV-10xHis vectors, resulting in the pYeDP60-PcCdc50B-3C_protease-GFP-10xHis vector.

### Yeast transformation

The different proteins and protein complexes were expressed in the *Saccharomyces cerevisiae* strain W3031b Gal4-2 (a, leu2, his3, trp1::TRP1-GAL10-GAL4, ura3, ade2-1, canr, cir+) (Lenoir et al. 2002). Yeast cells were transformed using the lithium acetate method (Gietz et al. 1995). 5 ml of non-transformed W3031b Gal4-2 strain was cultured in S6AU minimal media (0.1 % (w/v) Bacto™ Casamino Acids, 0.7 % (w/v) Yeast nitrogen base (without amino acids and with ammonium sulphate), 2 % (w/v) glucose, 20 μg/mL adenine and 20 μg/mL uracil) for 24 h at 28°C. Cells were mixed with ~1 μg of plasmid DNA and 100 μg of salmon sperm DNA (denatured at 100°C for 5 min and cooled down on ice). After vortexing, 500 μL of 40 % (w/v) PEG 4000, 100 mM lithium acetate, 10 mM Tris-HCl pH 7.5 and 1 mM EDTA and 20 μL of 1 M DTT was added. The tube was vortexed again and left overnight at room temperature. The next morning the tube was centrifuged at 350 *x g* for 2 min, the supernatant was removed, and the cells were re-suspended in 100 μL of S6A minimal media (SAU without uracil) and plated on S6A-agar plates. The plates were left in the incubator at 28°C for 3 to 5 days until the transformed colonies appeared.

### Protein expression in *S. cerevisiae*

Transformed yeast colonies were pre-cultured in 5 mL of S6A media or 24 h at 28°C. For small-scale expression tests, the pre-culture was diluted to OD_600_ of 0.2 in 20 mL of YPGE2X rich media (2% (w/v) Bacto™ Peptone, 2% (w/v), yeast extract, 1% (w/v) glucose and 2.7% (w/v) ethanol) and cultured for 30 h at 28°C. Then, the cell culture was cooled down to 18°C, and 20 g/L of galactose was added to induce protein expression for 18 h at 18°C.

For large-scale cultures used for membrane fractionation, the first pre-culture was diluted to a OD_600_ of 0.1 in 50 mL of S6A media and incubated at 28°C for another 24 h. Then, this second pre-culture was diluted to an OD_600_ of 0.05 in 500 mL of YPGE2X media. The cells were grown for 36 h at 28°C to consume the majority of glucose. Then, the culture flasks were cooled down to 18°C, and 2% (w/v) of galactose were added to induce protein expression. After 13 h at 18°C, a second addition of 2 % (w/v) galactose was done and the culture was left for five more hours. The cells were centrifuged at 4000 x *g* during 10 min at 4°C and washed two times with cold water. The cell-pellet was weighted and re-suspended in 2 mL of cold TEKS buffer (50 mM Tris-HCl pH 7.5, 1 mM EDTA, 0.1 M KCl and 0.6 M sorbitol) per gram of cells. After incubation at 4°C for 15 min, the cells were centrifuged and the resulting pellet was flash-frozen in liquid nitrogen and stored at −80°C.

### Membrane preparation

To analyze the expression of the proteins induced in the small-scale cultures, a total membrane preparation was done. A volume of each culture corresponding to 10 optical density units at 600 nm was centrifuged at 800 x *g* for 10 min and 4°C. Cells were re-suspended in 1 mL of ice-cold TEPI buffer (50 mM Tris-HCl pH 6.8, 5 mM EDTA, 20 mM NaN3) supplemented with SigmaFast™ EDTA-free Protease Inhibitor Cocktail (PIC) tablets (Sigma-Aldrich) and 1 mM phenylmethylsulfonyl fluoride (PMSF, Sigma-Aldrich) and transferred into 1.5 mL Eppendorf tubes. After centrifugation at 1000 x *g* for 10 min and 4°C, the pellet was re-suspended in 100 μL of ice-cold TEPI buffer, and cells were broken after adding 100 μL of glass beads (0.5 mm of diameter) and vortexing for 25 min at 4°C. Then, TEPI buffer was added to a final volume of 1 mL and samples were centrifuged for 5 min at 500 x *g* and 4°C. The supernatant was then centrifuged at 100 000 x *g* for 90 min and 4°C. The resulting membrane pellet was re-suspended directly in 200 μL of SDS-PAGE loading buffer and subjected to SDS-PAGE and western blot analysis.

Frozen cell-pellets from large-scale cultures were resuspended in 1 mL of TES buffer (50 mM Tris-HCl pH 7.5, 1 mM EDTA and 0.6 M sorbitol) per gram of cell-pellet, supplemented with PIC and 1 mM PMSF. The cells were broken with 0.5 mm glass beads using the planetary mill Pulverisette 6 (Fritsch). The broken crude extract was recovered and the beads were washed with 1.5 mL of ice-cold TES buffer per gram of cell-pellet. The pH of the broken cells was adjusted to 7.5, and centrifuged at 1000 x *g* for 20 min at 4°C. The resulting supernatant (S1) was centrifuged at 20 000 x *g* for 20 min at 4°C to obtain the second pellet (P2) or high-density membrane fraction. The resulting supernatant from the previous step (S2) was then centrifuged at 125 000 x *g* during 1 h at 4°C, obtaining the third pellet (P3) or low-density membrane fraction. P3 pellet was re-suspended in 0.2 mL of HEPES-sucrose buffer (20 mM HEPES pH 7.6, 0,25 M sucrose) per gram of cell-pellet. Aliquots of all membrane pellets were flash-frozen in liquid nitrogen and stored at −80°C.

### Protein detection by western blotting

Total protein concentration in the different samples was measured by the Bicinchoninic Acid Assay (BCA) (Walker 2003). Samples were subjected to SDS-PAGE and proteins were transferred to Polyvinylidene difluoride (PVDF) membranes (Immobilon^®^-P, Merck). The ATP2 proteins fused to the BAD tag were detected with the Avidin HRP-conjugate probe (Invitrogen), diluted to 1:20 000 in 5 % (w/v) milk in PBS-Tween buffer (PBS + 0.02 % (v/v) Tween^®^20). Cdc50 proteins fused to the 10xHis tag were detected using the HisProbeTM-HRP conjugate (ThermoFisher) diluted to 1:2000 in 2 % (w/v) BSA in PBS-Tween buffer. GFP-tagged proteins were detected with a primary mouse antibody IgG1K Anti-GFP (Roche) diluted to 1:1000 in 5 % (w/v) milk in PBS-Tween buffer, followed by a incubation with a second Goat Anti-Mouse IgG-HRP conjugate antibody (Bio-Rad) diluted to 1: 3000 in 5 % (w/v) milk in PBS-Tween buffer. The ECL Western Blotting Detection kit (GE healthcare) was used for western blot revelation, and luminescence was detected by a CDD camera.

### Deglycosylation assays of PcCdc50 proteins

P3 membranes co-expressing PcATP2/PcCdc50A, PcATP2/PcCdc50B or Drs2p/Cdc50p were diluted to 4 mg/mL in HEPES-sucrose buffer. 20 μg of total protein were added to a glycoprotein denaturing buffer (0.5 % (w/v) SDS, 40 mM DTT) and protein denaturation was performed at 100°C for 10 min in a dry bath. Samples were cooled down on ice for 5 min and spun down at 15 000 *x g* for 10 sec. 10 μL of denatured samples were enzymatically digested with either, PGNase F or EndoH (New England Biolabs) using the buffers and protocols provided by the supplier. The reaction was carried out for 1 h at 37°C and results were analyzed by western blot.

### Detergent solubilization screenings

P2 and P3 membrane fractions were solubilized with eleven different detergents at 2 mg/mL of total protein concentration in the membranes, and 10 mg/mL of detergent concentration in solubilization buffer (20 mM Tris-HCl pH 7.8, 10 % (v/v) glycerol, 150 mM NaCl, 1 mM PMSF and PIC). Solubilization was performed at 4°C or 20°C and at different incubation times. 1 h and 20°C were the conditions that showed the best results. After incubation, samples were ultracentrifugated for 1 h at 125 000 x *g* and 4°C, and both the pellet and the supernatant (detergent-solubilized sample) were subjected to SDS-PAGE and western-blot to analyze the solubilization efficiency of each experimental condition. Detergents screened: n-decyl-β-D-maltopyranoside (DM), n-undecyl-β-D-maltopyranoside (UDM) and octaethylene glycol monododecyl Ether (C12E8) from Calbiochem; n-dodecyl-β-D-maltopyranoside (DDM), lauryl maltose neopentyl glycol (LMNG), and n-dodecyl phosphocholine 12 (FosC12) from Anatrace; n-Octyl-β-D-glucopyranoside (OG), n-Octyl-β-D-thioglucopyranoside (OTG) from Sigma-Aldrich; and Laurydimethylaminoxide (LDAO), and 5-cyclohexyl-1-pentyl-β-D-maltoside (CYMAL-5) from Fluka. Solubilization experiments were also performed in the presence of the cholesterol derivative, cholesteryl hemisuccinate (CHS) (Sigma-Aldrich) at a 5:1 (w/w) detergent to CHS ratio.

### Co-immunoprecipitation assays

Co-immunoprecipitation assays were performed using agarose beads coupled to nanobodies recognizing the GFP (GFP-Trap^®^, ChromoTek), and following the protocol suggested by the provider. 500 μL of P3 membranes diluted to 2 mg/mL of total protein concentration were solubilized in solubilization buffer containing 10 mg/mL of DDM and 2 mg/mL CHS, during 1 h at 20°C. After incubation, membranes were ultra-centrifuged at 125 000 x *g* for 90 min at 4°C, and the supernatant was incubated for 1 h and 4°C with gentle rotation with 25 μL of GFP-Trap^®^ beads, previously equilibrated in washing buffer 1 (20 mM Tris-HCl pH 7.8, 150 mM NaCl, 0.2 mg/mL DDM, 0.04 mg/mL CHS, 10 % (v/v) glycerol). After incubation, the flow-through was removed by centrifuging the sample at 100 x *g* for 30 sec. The beads were then washed by adding 500 μL of ice-cold washing buffer 1 followed by centrifugation at 100 x *g* for 30 sec. A second wash was done with 500 μL of washing buffer 2 (20 mM Tris-HCl pH 7.8, 500 mM NaCl, 0.2 mg/mL DDM and 10 % (v/v) glycerol). Proteins bound to the GFP-Trap^®^ beads were eluted by adding 50 μL of 0.2 M glycine buffer pH 2.5 followed by centrifugation at 100 x *g* for 30 sec. The low-pH of the elution was quickly neutralized by adding 5 μL of 1 M Tris-base pH 10.4. This elution step was repeated one more time and samples were analyzed by western blot.

### Fluorescent size exclusion chromatography

P3 membranes were solubilized as described in the previous section and 200 μL of the detergent-solubilized supernatant was injected in a Superose 6 10/300 GL gel-filtration column (GE healthcare) equilibrated with 20 mM Tris-HCl pH 7.8, 150 mM NaCl, 10 % (v/v) Glycerol, 0.1 mg/mL DDM, 0.02 mg/mL CHS. Fluorescent size-exclusion chromatography (FSEC) experiments were performed in an ÄKTA™ purifier chromatography system (GE healthcare) with a fluorescence detector (FP-4025, JASCO) connected “in line” with the column. The excitation and emission wavelengths of the fluorescence detector were set, respectively, to 460 and 520 nm. Due to the approximately 10 times difference on expression yield between PcCdc50B-GFP and PcATP2-GFP, the gain (or output sensitivity) of the fluorescence detector was set to x 100 in the experiments were the GFP was fused to PcATP2 and x 10 in the experiments were the GFP was fused to PcCdc50B. FSEC chromatograms were plotted using Plot2 software and data was normalized.

### Generation of home-made GFP-Trap^®^ beads for ATPase assays

We first expressed and purified a nanobody directed against GFP (nanoGFP). The plasmid vector pOPINE GFP nanobody was a gift from Brett Collins (Addgene plasmid # 49172) (Kubala et al. 2010). 500 mL of *E. coli* BL21 cells harboring this plasmid were cultured at 37°C in LB media supplemented with ampicillin. When the OD_600_ reached ~0.6, the temperature was dropped to 20°C, and protein expression was induced with 1 mM isopropyl β-D-1-thiogalactopyranoside (IPTG) (Sigma-Aldrich) for 20-24h at 20°C. The cells were spun down, flash-frozen in liquid nitrogen, and stored at −80°C. The cell-pellet was re-suspended in 10 mL of lysis buffer (PBS pH 8, 0.5 M NaCl, 5 mM imidazole, 1 mM PMSF and 10 μg/μL of Lysozyme) and left for 1 h at 4°C with gentle rotation. Cells were broken by sonication, maintaining the cells always on ice. Cell lysate was centrifuged at 20 000 x *g* for 20 min at 4°C, and the supernatant was mixed with 1 mL of TALON^®^ metal affinity beads (Clontech) previously equilibrated with equilibration buffer (PBS pH 8, 0.5 M NaCl and 5 mM imidazole), and incubated for 1 h at 4°C under gentle rotation. After incubation, the beads were sequentially washed with 20 column-volumes of equilibration buffer, 10 column-volumes of equilibration buffer with 20 mM imidazole, and 10 column-volumes of equilibration buffer with 30 mM imidazole. The nanoGFP was eluted with two steps of 5 column-volumes of equilibration buffer with 150 mM of imidazole, and 5 column-volumes of equilibration buffer with 300 mM imidazole. The elution fractions were analyzed by SDS-PAGE, pooled and concentrated using a Vivaspin^®^ 20 5000 Mw concentrator (Sartorius). The concentrated protein was dialyzed overnight at 4°C to remove the imidazole, further concentrated and stored at 4°C until use. The purified nanoGFP was covalently bound to agarose beads following a previous protocol (Rothbauer et al. 2008). 1 ml of NHS-Activated Sepharose 4 Fast Flow beads (GE Healthcare Life Sciences) was washed with 10-15 column-volumes of 1 mM HCl, and equilibrated with PBS pH 7.5. Then, 1 mg of nanoGFP was added to the beads and left overnight at 4°C under gentle rotation, keeping a nanoGFP to beads volume ratio of approximately 0.5 to 1. After reaction, the remaining non-reacted sites were blocked by the addition of 10-15 column-volumes of 0.1 M Tris-HCl pH 8.5 followed by a 4 h incubation with this buffer at 4°C. Finally, the beads were subjected to 3 cycles of two sequential washes of 3 column-volumes of 0.1 M Tris-HCl, 0.5 M NaCl, pH 8.5 and 3 column-volumes of 0.1 M Acetate buffer, pH 5.0, 0.5 M NaCl. Finally, the home-made GFP-Trap^®^ beads were equilibrated in PBS pH 8.0 and stored at 4°C until use.

### Purification and immobilization of PcATP2/PcCdc50B in home-made GFP-Trap^®^ beads

P3 membranes expressing either PcATP2-GFP-BAD/PcCdc50B-His or PcATP2-D596N-GFP were diluted up to 5 mg/mL of total protein concentration in solubilization buffer (20 mM Tris-HCl pH 7.8, 150 mM NaCl, 100 mM KCl, 5 mM MgCl2, 20 % (w/v) glycerol, 20 mg/mL DDM and 4 mg/mL CHS), supplemented with PIC and 1 mM PMSF. After 1 h at 20°C under gentle rotation, samples were ultracentrifuged at 125 000 x *g* during 1 h at 4°C, and the supernatant (detergent-solubilized proteins) were mixed with the home-made GFP-Trap^®^ beads previously equilibrated with 10 column-volumes of washing buffer (20 mM Tris-HCl pH 7.8, 150 mM NaCl, 100 mM KCl, 5 mM MgCl2, 20 % (w/v) glycerol, 0.5 mg/mL DDM, 0.1 mg/mL CHS and 0.01 mg/mL POPS). After 2 h at 4°C of incubation with gentle rotation, the beads were washed with 20 column-volumes of washing buffer. GFP-Trap^®^ beads with bound PcATP2-GFP-BAD/Cdc50B-10xHis were diluted twice in washing buffer and flash-frozen in liquid-N2. The concentration of PcATP2-GFP-BAD in the beads was quantified by measuring the GFP fluorescence in a 96-well plate, and using a standard-calibration curve of purified GFP as reference. The protein content coupled to the GFP-Trap^®^ beads was eluted with 0.2 M glycine buffer pH 2.5 prior analysis by Coomassie-blue staining and western blot.

### ATPase assays

Between 5 and 10 μL of home-made GFP-Trap^®^ beads coupled with PcATP2-GFP-BAD/Cdc50B-10xHis (wild-type and mutant), and corresponding to 100 ng of PcATP2-GFP-BAD, were diluted into 50 μL of the reaction buffer (50 mM MOPS-Tris pH 7.0, 100 mM KCl, 5 mM MgCl2, 20 % (w/v) glycerol, 1 mM NaN3, 20 mg/mL DDM, 4 mg/mL CHS), supplemented with 0.45 mg/ml of the phospholipid substrate and, when indicated, with 0.025 mg/ml of PI4P. Samples were incubated on ice for 15 minutes. The reaction was initiated after addition of 1 mM ATP-Mg^2+^ and incubated at 37°C for 25 min. The reaction was stopped by dropping the test tubes into liquid N2. To quantify the hydrolyzed ATP, we used the BIOMOL^®^ Green solution (Enzo Biochem Inc), a proprietary mixing solution that uses a molybdate/malachite green-based assay to quantify the released Pi (Chan, Delfert, and Junger 1986; Hiraizumi et al. 2019). Thus, 100 μL of BIOMOL^®^ Green solution was added to each sample, vortexed and incubated at room temperature for 30 min. After incubation, samples were spun down to pellet the beads, and 100 μL of clear supernatant were deposited in a 96-well plate to measure the absorbance at 620 nm. Standard calibration curves at each experimental buffer condition were made using a phosphate standard solution provided by the BIOMOL^®^ Green supplier.

### PcATP2 3D model

To build the PcATP2 model, a divide and conquer strategy was taken. TOPCONS algorithm (Tsirigos et al. 2015) was used to infer membrane protein topology, resulting in 10 transmembrane (TMD) segments. The large intracellular domains (ICD) between TM2 and TM3 (ICD1) and TM4 and TM5 (ICD2) were aligned using MAFFT (Katoh et al. 2002) and modeled independently in MODELLER (Fiser and Šali 2003). ICD1 was modeled against templates 6KG7, 6ROH, and 1IWO. ICD2 was modeled against templates 6K7G, 6ROH, 1IWO, 4HQJ, and 5MPM. The TM domain (TMD) was modeled by carbon alpha molecular replacement in UCSF Chimera (Pettersen et al. 2004) using 6K7G and 6ROH as templates. The whole PcATP2 sequence was realigned using MAFFT and manually refined against the ICD1, ICD2, and TMD. This alignment was used as template to build the overall PcATP2 model in MODELLER. PcATP2 model was embedded in an POPC bilayer, energy minimized, and equilibrated for 10 ns using the CHARMM36F (Feller and MacKerell 2000) force field under NPT conditions, as described previously (L.-Y. Wang et al. 2016).

