## Supplementary material for "ATP2, the essential P4-ATPase of malaria parasites, catalyzes lipid-dependent ATP hydrolysis in complex with a Cdc50 β-subunit": Figures supplement

[1]Université Paris-Saclay, CEA, CNRS, Institute for Integrative Biology of the Cell (I2BC), 91198, Gif-sur-Yvette, France, CEA and Institut des Sciences du Vivant FREDERIC-JOLIOT, 91191, Gif-sur-Yvette, France. [2]Department of Medical Biochemistry and Biophysics, Umeå University, Umeå, Sweden. [3] Department of Clinical Medicine (FMUSP), University of São Paulo São Paulo, Brazil.[4]Biophysics Unit, Department of Biochemistry and Molecular Biology, School of Medicine, Universitat Autònoma de Barcelona, 08193 Cerdanyola del Vallés, Catalonia, Spain.

[*]Correspondance to : José Luis Vázquez-Ibar:.

### SUPPLEMENTARY FIGURES LEGENDS

#### Figure supplement 1

**Multiple sequence alignments of ATP2 homologs encoded by Apicomplexan parasites**. The figure only shows the regions where conserved residues or motifs mentioned in the main text are located. Lines above define the predicted transmembrane domains of PfATP2 (PF3D7_1219600) calculated using TOPCONS (Tsirigos et al. 2015). Conserved regions of these sequences specific to P-type ATPases or P4-ATPases are framed. Residues involved in the coordination of the PS head group in the human ATP8A1 (Hiraizumi et al. 2019) are highlighted with a grey background. Protein codes used by the Eukaryotic Pathogen Database Resources (EuPathDB): **PF3D7_1219600**, *P. falciparum* ATP2 (PfATP2); **PVX_123625**, *P. vivax* ATP2; **FBANKA_1434800**, *P. berghei* ATP2; **PCHAS_1436800**, *P. chabaudi* ATP2 (named PcATP2 in this work); **TGME49_247690**, *Toxoplasma gondii* ATP2 homolog; **cgd7_1760**, *Cryptosporidium parvum* ATP2 homolog, and **BMR1_01G01915** *Babesia microti* ATP2 homolog. (*) indicates positions which have a single, fully conserved residue, (:) indicates conserved residues of strongly similar properties, and (.) indicates conserved residues of weakly similar properties.

#### Figure supplement 2

**Multiple sequence alignments of *Plasmodium* and Apicomplexan Cdc50B homologs.** The figure only shows the regions where conserved residues or motifs mentioned in the main text are located. Lines above define the predicted transmembrane domains (TMDs) of PfCdc50B (PF3D7_1133300) calculated using TOPCONS (Tsirigos et al. 2015). The residues highlighted with a grey background correspond to the four conserved cysteines involved in the formation of the two disulfide bridges of the Cdc50 ectodomain. Potential residues in PcCdc50B (PCHAS_093090) involved in O-glycosylation identified by NetOGlyc - 4.0 are framed. Protein codes used by the Eukaryotic Pathogen Database Resources (EuPathDB): **PF3D7_1133300** *P. falciparum* Cdc50B (PfCdc50B); **PVX_092270**, *P. vivax* Cdc50B; **PBANKA_091510** *P. berghei* Cdc50B; **PCHAS_093090**, *P. chabaudi* Cdc50B (named PcCdc50B in this work); **TGME49_230820,** *Toxoplasma gondii* Cdc50B homolog; **cgd5_**360, *Cryptosporidium parvum* Cdc50B homolog, and **BMR1_03g01157**, *Babesia microti* Cdc50B homolog. (*) indicates positions which have a single, fully conserved residue, (:) indicates conserved residues of strongly similar properties, and (.) indicates conserved residues of weakly similar properties.

#### Figure supplement 3

**Detergent solubilization of P3 membranes co-expressing PcATP2 with either PcCdc50A or PcCdc50B.** Membranes co-expressing BAD-PcATP2/PcCdc50B-His (panels *A* to *D*) or BAD-PcATP2/PcCdc50A-His (panels *E* and *F*) at 2 mg/ml of total protein concentration were solubilized in 1 % (w/v) of the indicated detergent for 1 h at 20°C, in the presence or absence of 0.2 % (w/v) of cholesteryl hemisuccinate (CHS). After ultracentrifugation, 1 µg of total protein of the pellet (*P*, non-soluble material) and the supernatant (*S*, soluble material) were loaded on each line. Panels *A*, *C* and *E*, western blots revealed with the probe against the BAD. Panels *B*, *D* and *F*, western blots revealed with the HisProbe™ to detect the 10xHis tag. *DDM*, N-dodecyl-β-D-maltopyranoside, *LMNG*: Lauryl maltose neopentyl-glycol, C12E8, Octaethylene glycol monodecyl ether.

**Figure supplement 4**

**Western blot analysis of P3 membranes co-expressing PcATP2-GFP-BAD (wild-type and D596N mutant) with PcCdc50B-His, and PcATP2-GFP-BAD with PcCdc50A-His.** Line 1, co-expression of PcATP2-GFP-BAD with PcCdc50A-His; Line 2, co-expression of PcATP2-GFP-BAD with PcCdc50B-His; Lane 3, co-expression of D596N-PcATP2-GFP-BAD with PcCdc50B-His. Top panels, western blots revealed with an antibody against the GFP to detect PcATP2-GFP-BAD. Bottom panels, western blots revealed with the HisProbe™ to detect the 10xHis tag of PcCdc50 proteins.

**Figure supplement 5**

**Conservation in *Plasmodium* ATP2 orthologs of the PI4P binding-site and the autoinhibitory domains of Drs2p**. The figure shows the partial sequences alignments of *Plasmodium* ATP2 sequences and Drs2p. Lines above define the predicted transmembrane domains of PfATP2 (PF3D7_1219600) calculated using TOPCONS (Tsirigos et al. 2015). Residues at the C-terminal end involved in the binding of PI4P in Drs2p (Timcenko et al. 2019), and also conserved in the *Plasmodium* sequences are highlighted with a grey background. Protein motifs in Drs2p involved in protein autoinhibition are framed (Timcenko et al. 2019). Protein codes used by the Eukaryotic Pathogen Database Resources (EuPathDB): **PF3D7_1219600**, *P. falciparum* ATP2 (PfATP2); **PVX_123625**, *P. vivax* ATP2; **PBANKA_1434800**, *P. berghei* ATP2; **PCHAS_1436800**, *P. chabaudi* ATP2 (named PcATP2 in this work). Drs2p is a P4-ATPase of *S. cerevisiae*. (*) indicates positions which have a single, fully conserved residue, (:) indicates conserved residues of strongly similar properties, and (.) indicates conserved residues of weakly similar properties.

**REFERENCES FROM SUPPLEMENTARY INFORMATION**


Hiraizumi, Masahiro, Keitaro Yamashita, Tomohiro Nishizawa, and Osamu Nureki. 2019. "Cryo-EM Structures Capture the Transport Cycle of the P4-ATPase Flippase." *Science* 365 (6458): 1149–55. https://doi.org/10.1126/science.aay3353.

Timcenko, Milena, Joseph A Lyons, Dovile Januliene, Jakob J Ulstrup, Thibaud Dieudonné, Cédric Montigny, Miriam-Rose Ash, et al. 2019. "Structure and Autoregulation of a P4-ATPase Lipid

Flippase." *Nature* 571 (7765): 366–70. https://doi.org/10.1038/s41586-019-1344-7.

Tsirigos, Konstantinos D, Christoph Peters, Nanjiang Shu, Lukas Käll, and Arne Elofsson. 2015. "The TOPCONS Web Server for Consensus Prediction of Membrane Protein Topology and Signal Peptides." *Nucleic Acids Research* 43 (W1): W401-7. https://doi.org/10.1093/nar/gkv485.

### Figure supplement 1

```
                          TMD1               TMD2
PfATP2         FHKISNVYFFFIGILQVIPQFTATNGIPTVFFPLLIVLTANAIKDAFEDWNRHKTDKIEN 116
PVX_123625     FHKISNVYFLIIGILQLVPEFTATNRLPTILFPLTIVLVANAIKDAYEDWNRHKTDKIEN 116
FBANKA_1434800 FHKISNIYFFIIGVLQLVPELTATNRIPTILFPLSIVLIANAINDAYEDWNRHRTDKIEN 109
PcATP2         FHKISNVYFFIIGVLQLVPELTATNRIPTILFPLSIVLIANAINDAYEDWNRHKTDKIEN 109
TGME49_247690  FHKVSNVYFVVICCLQMIPQISTTNGVPTLALPLSIVLVVNAAKDAFEDWQRHRSDRIEN 109
cgd7_1760      FCRPVNFYFLVISLLQIFPSISSTNGIPTLALPLVFVLFVGAVKDGWEDLNRHQNDRIEN 115
BMR1_01G01915  LKQPLSLYFLAIAILQITPSISATRGIPTVMLPLFIVIAIDSIKDAYEDWQRHTSDRAEN 112
               : :  ..**. *  **: *.:::*. :**: :** :*:  .: :*.:** :** .*: **
------------------------------------------------------------------------------
PfATP2         IAFVETSSLDGETNLKVKEANTFLFNILGNDRNSAIDNVKNLKGFILSDKPNKDLSTMYG 288
PVX_123625     IAFVETSSLDGETNLKVKEANGFVFNILTSDRGEAIEKVKNLKGFIISEKPNKDLTTMYG 287
FBANKA_1434800 ICFAETSSLDGETNLKVKEVNKYIFNNLTYNMDEAIEKAKKLRGYILSEKPNKNLSTMNG 288
PcATP2         ICFAETSSLDGETNLKVKEVNKYIFNNLTYNIDEAIEKVKKLRGYILSEKPNKNLSTMNG 283
TGME49_247690  GAFVETASLDGETNLKLKQTHRVTFEWLGSFLPLAVCYLLTRAGRIRCQVPNRDLNTYEG 273
cgd7_1760_     DVFIDTSSLDGESNLKRRFSHKESTKMLGNNIHDVIKRARYLEGLIECSPPGKDLHNFDG 276
BMR1_01G01915  ISYVETLCLDGETNLKRKEAVQITQNYLKNDLINVLERIKNCEASILCNVPDTDLDKFKG 251
                 : :* .****:*** :       : *      .:       . * .. *. :* .  *
------------------------------------------------------------------------------
                                         TMD4
PfATP2         PFNLE---------KAKKPYIVGIISFFSWVVITGNFVPISLIVTMSFVKVVQAYFISCD 518
PVX_123625     PFNLE---------ESKKPFIVGVISFFSWVVITANFIPISLIVTMSFVKVVQAYFISCD 496
FBANKA_1434800 PFNLV---------EPKAPIISGIVSFFSWIVITANFIPICLIVTMSFVKVVQAYFISCD 490
PcATP2         PFNLV---------EPKAPIVSGIISFFSWIVITANFIPICLIVTMSCVKVIQAYFISCD 480
TGME49_247690  -TAAG---------NSEGPVVFCILNFFTWMVLTCNLVPISLVLQMGMVKALQSLFIAQD 453
cgd7_1760_     GVSDVNEISYRATGQAIPISFVPVVRFCTWIVLLANIIPIALVVSMKIVKAIQGQFISRD 447
BMR1_01G01915  -SPFY---------KDVTEVRVVCTSFFTWISITCNVIPISAVVTMNLVRFIQGYFISVD 409
                             :           * :*: :  *.:**. :: *  *: :*. **: *

------------------------------------------------------------------------------
PfATP2         ELGQIEYIFSDKTGTLTCNIMEFRKCAINGISYGKGLTEIKRNILKKKNLEIPVEPTM-K 708
PVX_123625     ELGQIEYIFSDKTGTLTCNVMEFRKCAINGISYGNGLTEIKKHILKKKNMAIPEEPVL-K 684
FBANKA_1434800 ELGQIEYIFSDKTGTLTCNVMEFRKCAINGISYGTGLTEIKRKILKKNNIPIPQEPVDFD 656
PcATP2         ELGQIEYIFSDKTGTLTCNVMEFRKCAINGISYGTGLTEIKRKILIKNNIPIPPEPVDLD 645
TGME49_247690  ELGQVSYIFSDKTGTMTSNVMEFRKCCVRGLSYGQGLTEVRRQALRRLGLPVPADPLPPP 933
cgd7_1760_     DLGQVRYIFSDKTGTLTRNIMEFKSLSVGGVHYGSTETSSSKEDNLIREIEIPQ------ 522
BMR1_01G01915  LLGQVQICFSDKTGTLTCNKMNFRKFSIEGVSYGKGLTDIKRSYLIKNGIPVPGAISG-K 489
                ***:   *******:* * *:*:. .: *: **   *.  :       : :*
------------------------------------------------------------------------------
                            TMD6
PfATP2         QKIYFEFLLHLFNVLFTAIPVVIHAVLDQDISLNTAMEKPNLYKLGIHHYYFNIRTFISW 1359
PVX_123625     QKIYYEFLLHLHNILFTAVPVVAHAILDKDVSLNTALVTPSLYKLGIYHYYFNISTFVSW 1318
FBANKA_1434800 QKIYYEFLLHLYNVMFTSLPIVILAILDKDVSLNTALKNPCLYKLGIHNFYFNINKFISW 1321
PcATP2         QKIYFEILLHSYNVLFTSLPIIILAILDKDVSINTALKNPCLYKLGIHNFYFNINKFISW 1288
TGME49_247690  QKFYFEFLYQMYNVVFTAIPITLYGVFDQDVDKKLALKYPQLYRCGQIDLYLNLRVFLKW 1463
cgd7_1760_     TRLYFDYWYQVYNVILSSVPIVVVSVFDFDVTKSESLSKPHLYSFGPENKFLNTKICLIY 1155
BMR1_01G01915  QLLYNDMLQQLFNIFFTAIPSIIFGSIEQDVDRNVTFKYPQLYKLGHINFYMNMRAFLTW 1017
                 :* :   : .*:.::::*    . :: *:  . ::  * **  *  . ::*    : :
```

### Figure supplement 2

```
                                         TMD1
                             ________________________
PfCdc50B      ------ERVVGPVWINKYSSMIYFLMFLFILNLSVGILILILSSKYIECRIPYEYKG--E 90
PVX_092270    ------EKVIGPIWVPTYCSIIVFLLFLFFFNLLVGVAILIISSNYIECRVPYEYKS--Q 90
PBANKA_091510 ------EKIFGPVFVYKYSTLIAFFIFLFILNLSIGIAILYLSSQYIECKIPYEYKS--Q 93
PcCdc50B      ------EKVFGPVFVYKYSTSIVFFIFLFILNLSVGIAILYLSSQYIECKIPYEYKT--Q 93
TGME49_230820 --QVHQEAGNGMYPLWSAGVVLRLCLLGALFFVSVGAWLIFEDEQHVECKLNYAEKTLQE 221
cgd5_360      NKVI--NNIERWIPFYTPHYLILIYIFVGITFITVGIFLQIFSNNTIECIINYEDSPG-N 137
BMR1_03g01157 VKILEWDIRDGVYMQRRSAPILILFIFILAINICISSLLWTRKVNFVECEIPYHQQP--V 145
                    :              : : ::     : :.  :   . : :** : *  .

PfCdc50B      TFTKYSIVKVTPEQCKGQ----KNLKELNG-NINVHYEILGMQQNHYKFVSGMKKEQLNG 145
PVX_092270    AYTKYSIVKVTPEHCKGN----ENLKQLKG-PINIHYEIYGVQQNHYRFLTSFKKEQLRG 145
PBANKA_091510 PYTKYSIIKVTPEHCKGR----ENLKELKG-KINVHYEIYGVQQNHYSFMKSFNAEQIGG 148
PcCdc50B      PYTKYSIIKVTPEHCKGH----ENLKELKG-EINVHYEIYGVQQNHYSFMKSFNTKQLGG 148
TGME49_230820 GSSRYLLKGISSAHCTRE------VNELKGEEISVYAEMGHFFQNDAQVLWSRNDRQLAG 275
cgd5_360      GKVIDTIVEIKSEHCNPSMINGNELKYLKG-DFFIYYQLRNFYQNNKSFIFSRSDRQLSG 196
BMR1_03g01157 GNPTFVTIKVTHKECNKD----DKFALLEADDIFVYYKITNYPHLESSLSNGIVQEQLAG 201
                       :.  .*.        .  *:.  : :: ::    :  .  .  .   .*: *

PfCdc50B      NIFLKKEELEECYPLITFSEGKKKKKLLHPCGIFPWNVFTDSYIFYDKEPDEVPFP---T 202
PVX_092270    DLFLQEKELSECFPLITYEQSG-TRKILHPCGILQWNVFTDSYIFYDKEPDESPFP---T 201
PBANKA_091510 GIDVYKHDLNQCYPLITYFKDR-INKILHPCGILPWSVFTDNYIFYDKEPDDAPFP---D 204
PcCdc50B      KIFVSKDYLNHCYPLITYFKDR-INKILHPCGVLPWHVFTDNYIFYDKEPEDAPFP---D 204
TGME49_230820 KIFTDPKDVRECEPLATAVVGN-VTKVLHPCGALAWAVFTDKYQFLEGTPEGDNDQVPMK 334
cgd5_360      ELIYNEETLSDCYPVIKDKQ----GKIFYPCGVATLTIFNDTFTILDGQN---------D 243
BMR1_03g01157 NVISDSKQLHNCAPLDSIEHKG-VKKILHPCGIHAWNVFNDKIRFYRSSPTGS-LA---A 256
               :    . : .* *: .        *:::***     :*.*.  :

PfCdc50B      PLPLKQNVEEITI-KYYRQFYKNPSPQNVQLYKDHIYFWMEPDIQYERLQ-ENKETNEKL 260
PVX_092270    PLPLKQRAEDITI-KYYRKFFKNPTRDIINLHKKRIYFWMDEEVQLKILQ-EHAETNDKL 259
PBANKA_091510 PLPLNERVEDITI-KYFRKFFKNPHPENIKLYKDKVYFWMDAKTQSEALH-ENIVANEKL 262
PcCdc50B      PLPLKERVEDITI-KYFRKFFKNPHPETIDLYKDKVYFWMDRTTQSEALH-ENIVANEKL 262
TGME49_230820 PIPLNQTQAVLLHSWPWQDMYKNPPAEDRAAVLDKVYFWMSPVDNDDGEDMYKTREEARA 394
cgd5_360      PIEIDDSIDTITF-KSDQINYKNIPEHELLNH---------------------------- 274
BMR1_03g01157 SIEIDESVPTSAM-PLEIQHFKNPTQDIVDKHKQHTYFWMLPENEDSKEM-DDDEC--LA 312
               : :.:              :**   .

                                              TMD2
                                    _______________________
PfCdc50B      IND---TKIIVISTSQYYMRTF-LIGFIFIIISIIALILCIFYLIRMNKYENK------- 366
PVX_092270    ISD---TKIIVISNADFYFNTT-LIGIFFIITAVFALLLSLLYFIRMKKHQFK------- 365
PBANKA_091510 TAE---AKAIIITEANFYINNN-LIGIIFTIISIFSLILSILYYMRMKKHKFMRQI-DE- 373
PcCdc50B      TAE---AKAIIITEANFYINNN-LIGIIFIIISIFSLILSILYYMRMKKHKFMRRI-ED- 373
TGME49_230820 VSSWKGKKAIVLVQKSRFGGRSLFIGIAYLSFGCLLTM--LVFYMLWKKWQYRREGEEIR 509
cgd5_360      VHFFNGSKHIVISQSTIFGGKNPYFGILYIISGILFILLSIYYIIRNKFNTNILG--DFR 391
BMR1_03g01157 VAKFNGTKSIILSIPRWPYGSSLSLEILHLVFTILTLLFTVIYATRNTNSSTFLQMYHES 429
                     * *::            : : .     :  :  : :    .
```

**Figure supplement 3**

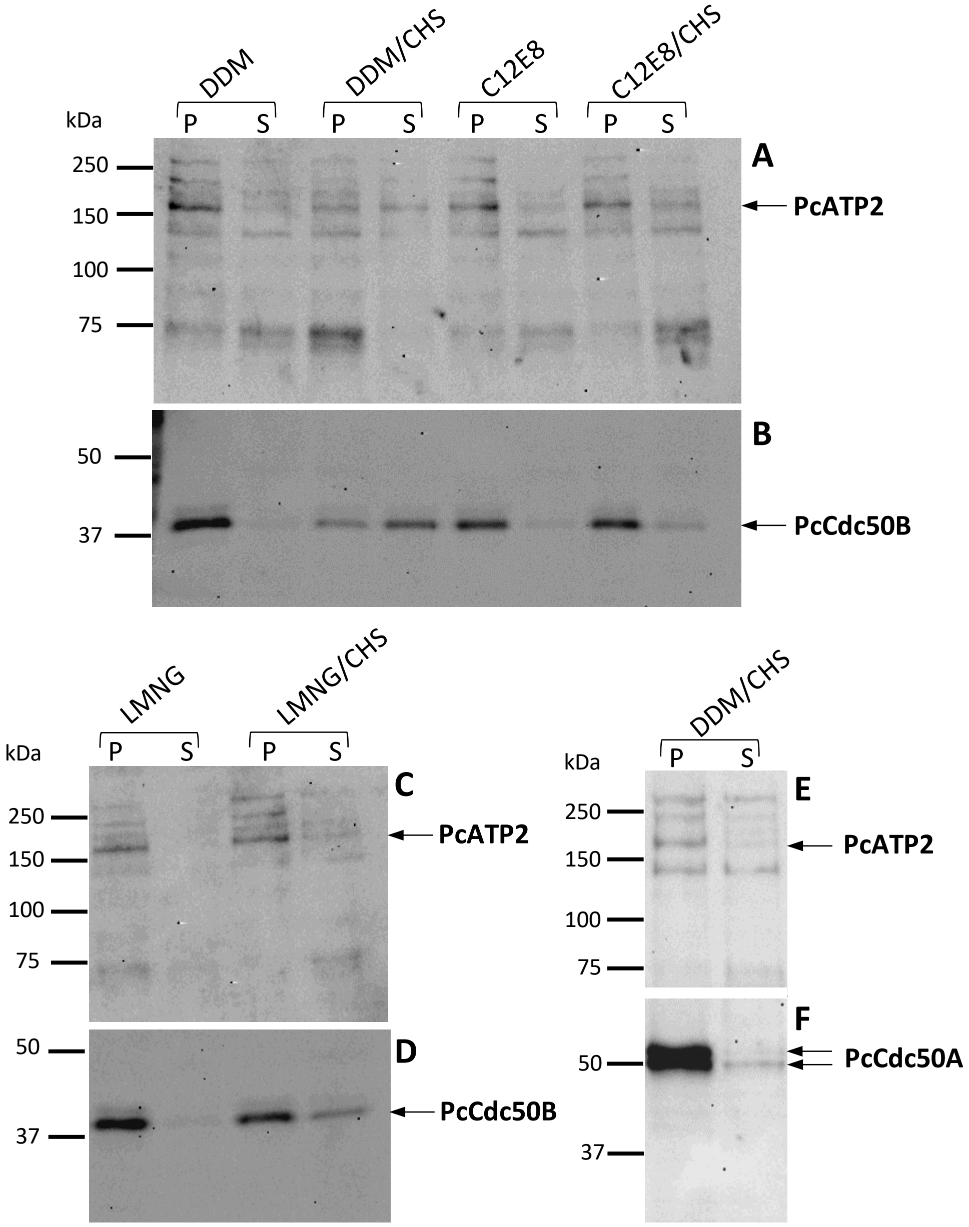

**Figure supplement 4**

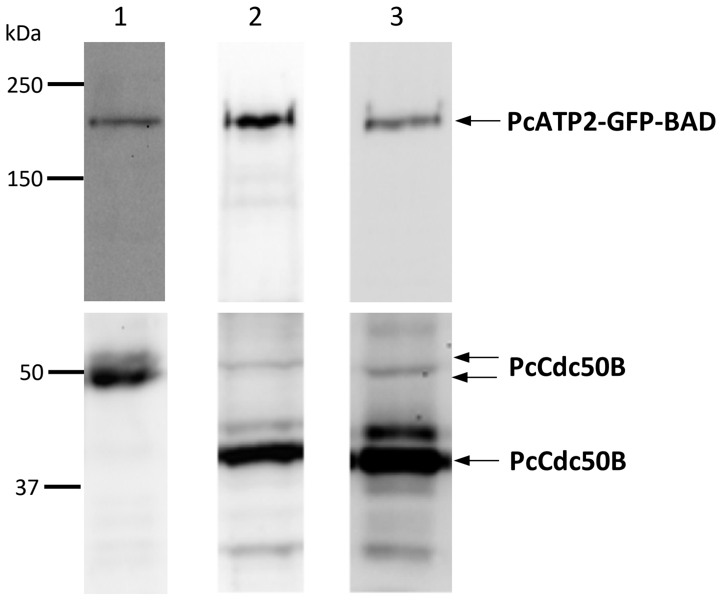

### Figure supplement 5

```
PfATP2         KHFDITFLYRREGKYGISIFG--KIYEIDTLATIEFTSKRKMSSVICRIPVINPDYNHPT 835
PVX_123625     KHFGITFLYRRDGKYGISIFG--TVYEIETLAIVEFTSKRKMSSVICRIPVRATHTGGAG 811
FBANKA_1434800 KHFGITFLYRKDGKCGIKIFD--KVYEIDILATVEFTSKRKMSTVVCRIPIISNESTEPS 782
PcATP2         KHFGITFLYRKDGKCGVKIFD--KVYEIDILATIEFTSKRKMSTIVCRIPVMSNEDTKTS 771
Drs2p          ADLGYKFIIRKPNSVTVLLEETGEEKEYQLLNICEFNSTRKRMSA--------------- 707
                .:. .*: *: ..  : :       * : *   **.*.**  :
-----------------------------------------------------------------------------
                      TMD10

PfATP2         RFWLVVILGLFTALLRDYVFKVYKRNFNPEIYHLLLDQENAKIGMNDVIDQLKLNEFDKD 1522
PVX_123625     RFWLVVLLGLFTALSRDFIFKVFKRNFNPEVYHFLLDQEDKPKGENNVINPLSSDPCQKE 1481
FBANKA_1434800 RFWLVLLLVLFTALTRDYVYKVYKKNFYPEAYHLLQDEEENISNTNHIQHS--SKCSNNM 1482
PcATP2         RFWLVFVLGLFAALTRDYVYKVYKKNFCPEAYHLLQDEEDKIENPKNIQHN--SRSSNNI 1449
Drs2p          VFWLTLIVLPIFALVRDFLWKYYKRMYEPETYHVIQEMQKYNISDSRPHVQQFQNAIRKV 1263
                ***..::  : ** **:::* :*: : ** **.: : :.   . .            :

PfATP2         DDIRIEKSKSLGYAFSEADPACIQLIRK-QDNMI-------------------------- 1555
PVX_123625     EEIKIEKCKSLGYAFSEVDPACVKLIRK-QDKLI-------------------------- 1514
FBANKA_1434800 DETKSSKSEFMGYAFSEADPACVHFIRK-QDKLI-------------------------- 1515
PcATP2         EEMKPSKSELMGYAFSEADPVCVNFIRK-QDKLI-------------------------- 1482
Drs2p          RQ-VQRMKKQRGFAFSQAEEGGQEKIVRMYDTTQKRGKYGELQDASANPFNDNNGLGSND 1322
                :      :  *:***:.:    . * :  *.
```
